# Predicting microbial growth conditions from amino acid composition

**DOI:** 10.1101/2024.03.22.586313

**Authors:** Tyler P. Barnum, Alexander Crits-Christoph, Michael Molla, Paul Carini, Henry H. Lee, Nili Ostrov

## Abstract

The ability to grow a microbe in the laboratory enables reproducible study and engineering of its genetics. Unfortunately, the majority of microbes in the tree of life remain uncultivated because of the effort required to identify culturing conditions. Predictions of viable growth conditions to guide experimental testing would be highly desirable. While carbon and energy sources can be computationally predicted with annotated genes, it is harder to predict other requirements for growth such as oxygen, temperature, salinity, and pH. Here, we developed genome-based computational models capable of predicting oxygen tolerance (92% balanced accuracy), optimum temperature (R^2^=0.73), salinity (R^2^=0.81) and pH (R^2^=0.48) for novel taxonomic microbial families without requiring functional gene annotations. Using growth conditions and genome sequences of 15,596 bacteria and archaea, we found that amino acid frequencies are predictive of growth requirements. As little as two amino acids can predict oxygen tolerance with 88% balanced accuracy. Using cellular localization of proteins to compute amino acid frequencies improved prediction of pH (R^2^ increase of 0.36). Because these models do not rely on the presence or absence of specific genes, they can be applied to incomplete genomes, requiring as little as 10% completeness. We applied our models to predict growth requirements for all 85,205 species of sequenced bacteria and archaea and found that uncultivated species are enriched in thermophiles, anaerobes, and acidophiles. Finally, we applied our models to 3,349 environmental samples with metagenome-assembled genomes and showed that individual microbes within a community have differing growth requirements. This work guides identification of growth constraints for laboratory cultivation of diverse microbes.

## Introduction

In order to grow a microorganism in the laboratory, its metabolic (nutrition and energy) and physicochemical (temperature, pH, salinity and oxygen) requirements must be met. Due to the vast combinatorial search space of possible chemical and physical parameters, robust conditions for culturing microbes are laborious to determine^1–3^. Computational predictions that could minimize the amount of experimental labor by constraining this search space would be highly desirable. Unlike metabolic requirements that can sometimes be predicted from known genes and pathways^4–8^, physicochemical requirements often lack known genetic determinants and are more difficult to predict.

Current methods to predict temperature, pH, salinity, and oxygen tolerance may not be broadly extensible to uncultivated microbes. Most microbes have only few experimentally characterized close relatives, limiting methods that rely on empirical data for phylogenetic relatives^9^. An alternative strategy is to correlate operational taxonomic units (OTUs) to physicochemical variables like pH and oxygen across habitats^10^. It is unclear if gene-based models can be more accurate^1,11–16^. First, the genes underlying these physicochemical adaptations in diverse organisms remain unknown. Second, most existing gene-based models for prediction of microbial growth conditions rely on phylogenetically-skewed databases for their training, leading to inaccurate predictions for new taxa^17,18^. To build predictive models for uncultivated microbes, careful consideration of phylogenetic bias is required. This includes training on phylogenetically balanced datasets, evaluation with phylogenetically novel examples (“out-of-clade prediction”), and using features that are less prone to spurious phylogenetic correlation^18^.

We hypothesized that models trained on features of DNA and protein sequences from microbes with known growth conditions could provide more robust and less phylogenetically-biased predictions for growth of uncultivated microbes. This is because changes in amino acid frequencies across an entire genome are less likely than losing or adding gene content, and thus more likely to be predictive across diverse organisms. For example, while some bacterial species can evolve to grow faster through the loss of many different genes^19^, maximum growth rate across diverse phyla is consistently correlated with bias in codon usage^20^. For oxygen sensitivity, aerobic organisms tend to have higher G+C content compared to anaerobic relatives, and amino acid frequencies have been linked to oxygen use^16,21–23^. For temperature, proteins tend to contain more aromatic and hydrophobic residues at higher temperatures^24–26^. For salinity, acidic amino acids and lower protein isoelectric points (pI) tend to be more frequent at higher salinity^27–29^. There have been no reported correlations of sequence composition with pH, possibly because the vast majority of proteins are intracellular and the cytoplasm remains close to neutral pH^30^.

Here, we evaluated the ability of models trained on properties of DNA and protein sequences to predict temperature, pH, salinity, and oxygen preferences of phylogenetically novel microorganisms (**Fig. 1A**). Models were trained on phylogenetically balanced data derived from 15,596 microbes and their performance was assessed using phylogenetically constrained out-of-clade testing. We then used the model to predict the cultivation requirements of 85,205 sequenced species of *Bacteria* and *Archaea* and metagenome-assembled genomes from 3,349 habitats with different physicochemical conditions.

**Figure 1.**
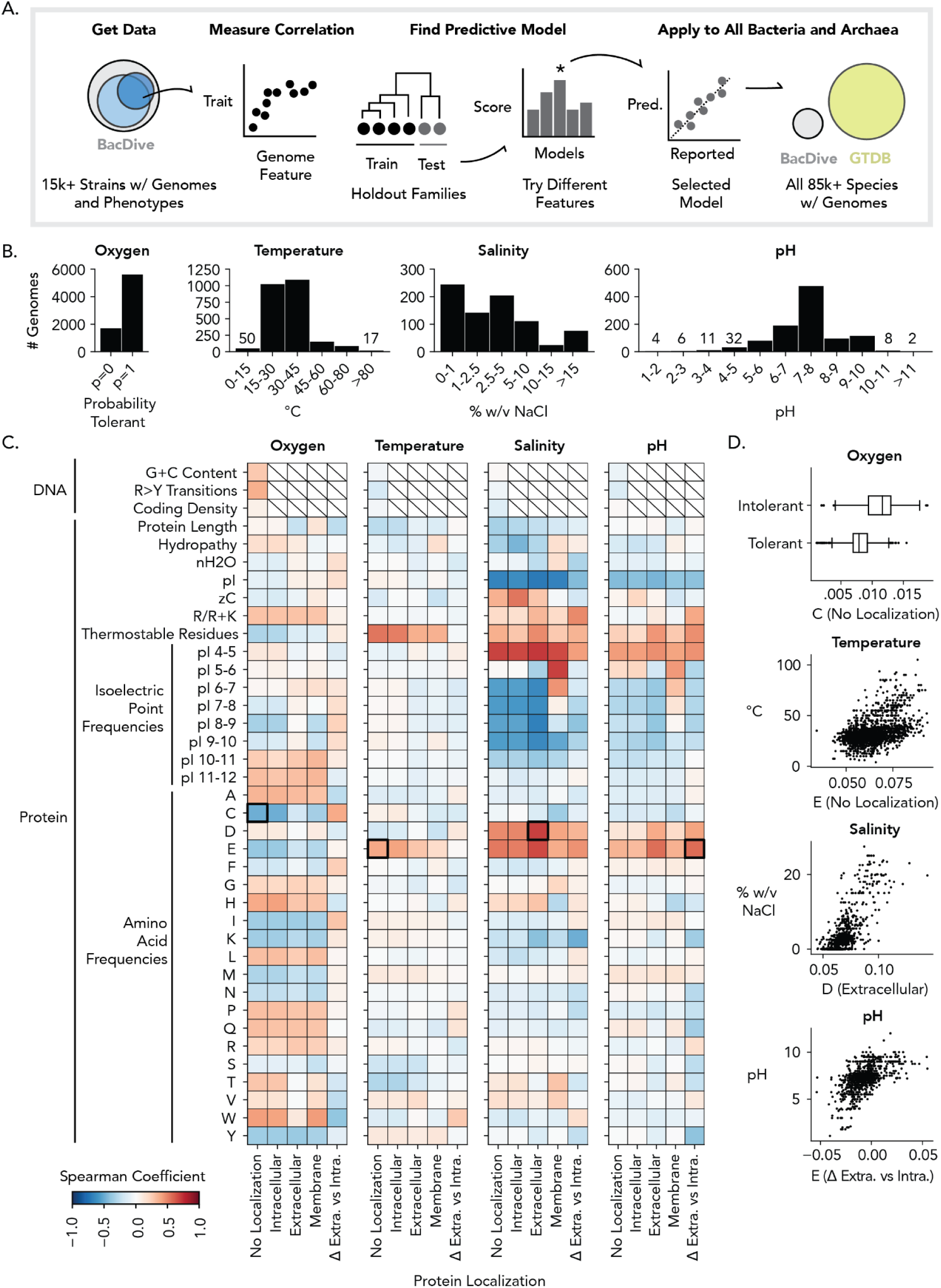
Overview and assessment of approach. **(A)** Schematic depiction of our approach. Only a fraction of organisms have reported growth conditions. Circles represent proportion between the number of all genomes in BacDive (gray), organisms in BacDive with empirical description of oxygen tolerance (dark blue), and organisms in BacDive with empirical description of temperature, salinity or pH (light blue). We hypothesized properties of DNA and protein sequences that can be used to predict temperature, pH, salinity, and oxygen. Our approach was to obtain and curate a dataset of growth conditions, explore their correlations to genome properties, and evaluate the ability of models to predict growth conditions for novel groups of organisms. Finally, we demonstrated potential applications of such models by expanding our prediction from the BacDive dataset (represented by gray circle) to all sequenced organisms (represented by yellow circle), totalling 85,205 prokaryotic genomes. Circle size is proportional to database size. **(B)** Distribution of curated data available for cultivated organisms as found in the BacDive dataset. **(C)** Correlation between physicochemical growth conditions and genomic features, as calculated for all genomes in the curated dataset. Growth conditions refer to optimum conditions unless otherwise noted. Each row represents one genome feature (refer to **Table 1** for feature descriptions). Color scale represents the degree of correlation: red indicates positive Spearman correlation coefficient, blue indicates negative). DNA sequence features were calculated for the whole genome. Protein sequence features were calculated based on predicted cellular localization of the resulting proteins. ‘Intracellular’ / ‘Extracellular’ / ‘Membrane’ represent values computed only on proteins with those localizations. ‘No localization’ represents calculation for all proteins regardless of predicted localization. ‘Δ Extra. vs. Intra.’ represents the difference between values computed on those sets. Black boxes indicate selected correlations shown in panel D. **(D)** Example of significant correlations observed between genetic features and growth conditions in panel C.

**Table 1.** Description of genomic variables used in this study. For proteins, variables were measured based on the localization of the protein to the intracellular, extracellular, or membrane compartments of the cell, the difference between extracellular and intracellular values, or based on no localization at all. References relate sequence features to growth conditions.

| Sequence feature | Variable name | Variable definition | Reference |
| --- | --- | --- | --- |
| DNA | G+C content | Fraction of bases that are guanine and cytosine | 23,24 |
|  | R>Y transitions | Fraction of bases that transition from a purine to pyrimidine | 24 |
|  | Coding density | Fraction of bases contained within coding sequences, approximated as the total length of proteins multiplied by three divided by genome length | 56 |
| Protein | Length | Mean number of amino acids across proteins |  |
|  | Hydropathy | Mean grand average of hydropathy (GRAVY) across proteins | 61 |
|  | nH<sub>2</sub>O | Mean stoichiometric hydration state, an estimate of the water molecules needed to compose a protein | 62 |
|  | Zc | Mean average carbon oxidation state, an estimate of the degree of oxidation of carbon in a protein | 62 |
|  | R/R+K | Ratio of arginine to arginine plus lysine | 54 |
|  | Thermostable residues | Sum of the amino acids I, V, Y, W, R, E, L reported to provide a correlation with high temperatures | 25 |
|  | pI | Mean isoelectric point across proteins, computationally estimated for each protein from the pKa of its amino acids | 27,28 |
|  | pI frequency | Fraction of proteins in the genome with a pI between assigned values, summing to 1 | 27,28 |
|  | Amino acid frequency | Frequency of each amino acid across all proteins | 22,25,28,54 |

## Results

### Overall model design

We chose to predict optimal growth conditions for four physicochemical factors: pH, temperature, salinity and oxygen tolerance (**Fig. 1A**). Importantly, we sought to build a predictive model without relying on gene functional annotations, where the only user input is a genome sequence.

First, we identified a set of DNA and protein sequence features that have been suggested to correlate with growth conditions (**Table 1**). For DNA sequences, we included G+C content (DNA 1-mers), frequency of purine-pyrimidine transitions, and estimated coding density. For protein sequences, we considered protein length, amino acid frequencies (amino acid 1-mers), predicted isoelectric point (pI), hydrophobicity, number of thermostable residues, hydration state (nH2O), average carbon oxidation state (Zc), and proportion of arginine to lysine residues (R/R+K). In addition, we considered the frequency of proteins within specific pI ranges (“isoelectric point frequencies”). We decided to restrict features to 1-mers because using longer k-mers of DNA or amino acid sequences can introduce spurious phylogenetic correlations due to their intrinsic phylogenetic signal^31^ and their high-dimensionality relative to the amount of training data available^18^.

To detect any potential influence of extracellular salinity and pH on protein composition, we chose to calculate protein features not only for all cellular proteins but also for subsets of intracellular-, extracellular-, and membrane-localized proteins. Extracellular proteins were identified with a hidden Markov model (see **Methods**) and the numerical difference between extra- and intra-cellular proteins (“Δ Extra vs. Intra”) were explored for prediction.

Next, we obtained empirical data describing abiotic conditions for 15,596 sequenced microbes from the BacDive database^1^. To utilize phenotypic data in our model we performed the following adjustments (see **Methods**): for temperature, pH, and salinity, we ignored qualitative parameters (*e.g.* “thermophile”) and filtered for precise measurements by only using continuous numerical parameters that have multiple measured values (*e.g.* a microbe with a 30°C optimum is ignored if no other temperatures were tested). We also chose to represent oxygen tolerance as probability, assigning ‘1’ for organisms described as ‘obligate aerobe’, ‘aerobe’, ‘facultative anaerobe’, or ‘facultative aerobe’ and ‘0’ to organisms described as ‘obligate anaerobes’ or ‘anaerobes’ without other labels. After curation, the number of bacteria and archaea available were 7293 for oxygen tolerance, 2418 for temperature, 801 for salinity, and 1020 for pH.

The distribution of BacDive data after curation is summarized for each growth condition in **Figure 1B**. Curation of quantitative values removed some bias, but distributions remain imbalanced: 77% of organisms were labeled oxygen tolerant, 87% of organisms are mesophiles (15-45°C), 74% have an optimum salinity between 0-5% w/v NaCl, and 65% have an optimum pH between 6 and 8. **Table 2** summarizes the number of genomes and the range of values that were used for each condition.

**Table 2.** Phenotypic data used in this study. Data was obtained from BacDive. The total number of organisms with optimum values reported is described for (1) all organisms with data and genomes in BacDive, (2) the subset of those organisms that remained after automated curation, and (3) the subset of those organisms after a balancing step. The range and mean values for the organisms used in modeling are provided.

| Growth condition | Organisms with data and genomes | Organisms after curation | Organisms used in modeling | Min-Max values in modeling (Mean) |
| --- | --- | --- | --- | --- |
| Oxygen tolerance | 7293 | 7293 | 3646 | 0-1 (0.65) probability tolerant |
| Temperature | 4773 | 2418 | 1722 | 4-105 (33) °C |
| Salinity | 2374 | 801 | 593 | 0.0-27.5 (4.3) % w/v NaCl |
| pH | 3545 | 1020 | 756 | 1.1-12.0 (7.2) pH |

### Correlation between sequence features and optimum physicochemical conditions

To assess the prospect for successful modeling, we calculated the above sequence features for each BacDive genome (**Supp. Data 1**), and measured their correlation with oxygen tolerance, temperature, salinity, and pH (**Fig. 1C**). No strong correlation was observed using DNA sequence features. We observed several strong correlations using protein sequence features. In several instances we observed stronger correlation when protein localization was considered, likely explained by differences between intracellular and extracellular salinity and pH. For example, optimum temperature correlated with the overall frequency of glutamic acid (Spearman correlation, ρ=0.39), optimum salinity correlated with the frequency of aspartic acid among extracellular proteins (ρ=0.69), optimum pH correlated with the difference between glutamic acid frequency for extracellular and intracellular proteins (ρ=0.56), and oxygen tolerance was correlated with low overall cysteine frequency (ρ=-0.49) (**Fig. 1D**). Remarkably, 69% of anaerobes but only 7% of oxygen tolerant microbes have more than 1.1% cysteine on average. Strong correlations were also observed between oxygen tolerance and the frequency of other amino acids. For example, across most protein localizations, the frequency of tryptophan, histidine, and alanine are positively correlated with oxygen tolerance, while frequency of cysteine, glutamic acid, and tyrosine are negatively correlated.

### Optimal physicochemical conditions can be predicted solely from sequence

We next developed models to predict growth conditions for novel taxa from sequence features. For each growth condition, we tried several estimators (untrained models) and nine different sets of features, grouped by feature type and protein localization. We assessed the performance of each model (an estimator trained on a set of features) through 5-fold cross-validation with holdouts at the family level (**Supp. Fig. 1**, **Supp. Data 2**, **Supp. Data 3**). We selected a single model for each growth condition based primarily on performance but also preferring simpler estimators, fewer features, and more interpretable features.

The accuracy of the selected models is summarized in **Figure 2A** and **Table 3**. Overall, we assessed oxygen tolerance to have the highest prediction accuracy (F1=0.94), followed by salinity (R^2^=0.81, RMSE=2.8% w/v NaCl) and temperature (R^2^=0.73, RMSE=6.5 °C). Optimal pH was least accurately predicted overall (R^2^=0.48, RMSE=1.1 pH). We found that amino acid frequencies alone were sufficient to provide all four predictions (**Fig. 2B**). Protein localization slightly increased the accuracy of predictions for optimum temperature and salinity (**Supp. Fig. 1B-C**) and was critical for predicting optimum pH (**Fig. 2B**). Notably, isoelectric point frequencies could accurately predict optimum salinity but not other conditions. In all models, minima and maxima were predicted with roughly the same accuracy as optima when the same estimator and features were used (**Supp. Fig. 2**). When challenged with a test dataset composed of families not in the training dataset, model performance was consistent with training and cross-validation data yet with lower accuracy (**Table 3**). This difference is due either to the smaller size and distinct composition of the dataset or to overfitting during model selection. We expect these models to be predictive on novel families to the degree of average error (RMSE) between the test and cross-validation data (**Fig. 2A**, **Table 3**).

**Figure 2.**
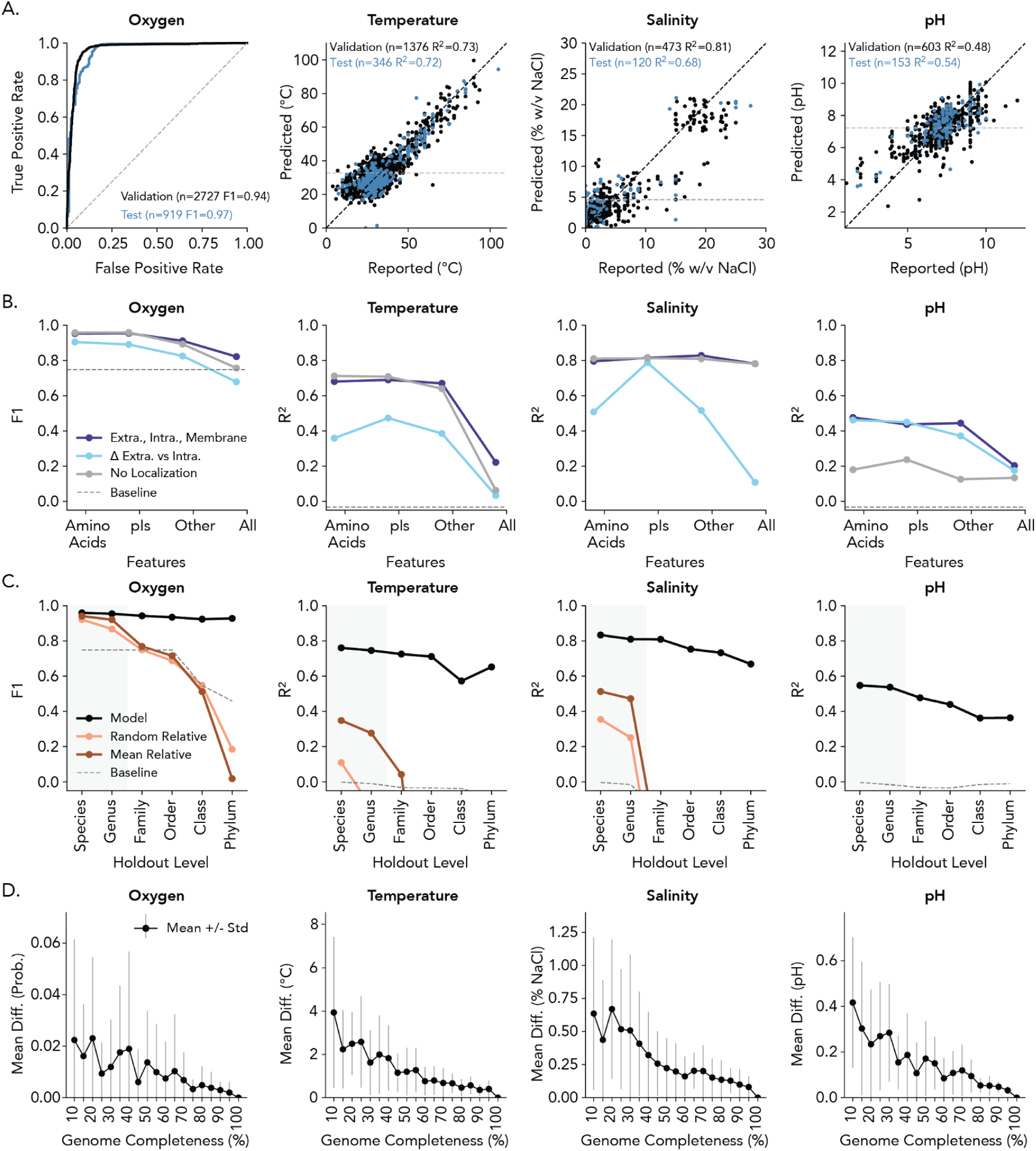
Accuracy of predictive models. **(A)** Accuracy of predictions for the selected model (estimator and features). Refer to **Table 3** for the estimator and features selected for each model. Prediction accuracy on validation (black) and test sets (blue) for optimum temperature, pH, and salinity are shown. Dashed gray lines indicate the baseline prediction, and solid gray lines indicate perfect predictions for regressions. Holdouts for the cross-validation and test datasets were performed at the family level, which means that the results reflect accuracy on new examples of families. **(B)** Performances during model selection when the estimator is held constant and the features are varied. Performance was evaluated in terms of coefficient of determination (R^2^) for regression or F1 score (F1) for classification. Negative R^2^ values, which result from random or otherwise less accurate prediction than selecting the mean, are not displayed. The types of features include: “AAs” amino acid frequencies; “pIs” isoelectric point frequencies; “Other” any feature not included in “AAs” or “pIs”; “All” all features. The localization of features can be: “No Localization” features were computed without considering localization; “Extra., Intra., Membrane” protein features were calculated on groups of proteins with the same localization; and “Δ Extra. Vs. Intra.” protein features were calculated as the difference between the values for extracellular and intracellular proteins. Dashed gray line represents the baseline for linear regression (always predicting the mean) or for logistic regression (always predicting the mode). **(C)** Influence of taxonomic holdout level on model performance. Black line represents the selected model evaluated using cross-validations performed at different taxonomic levels. Dark red line represents an alternative method provided for comparison, where the prediction is the average values of the closest relatives (*e.g.* when the holdout level is genus, the closest relatives are from other genera in the family at best). Light red line represents the same method but a random relative is chosen instead of using the average. Dashed gray line represents the baseline. Gray highlights indicate areas in which all methods tended to perform better than baseline. **(D)** Relationship between accuracy and genome completeness. Black dot is the mean error from 100% completeness and gray lines are ±1 standard deviation. Error was calculated using 20 genomes evenly distributed across percentiles of predicted values.

**Table 3.**
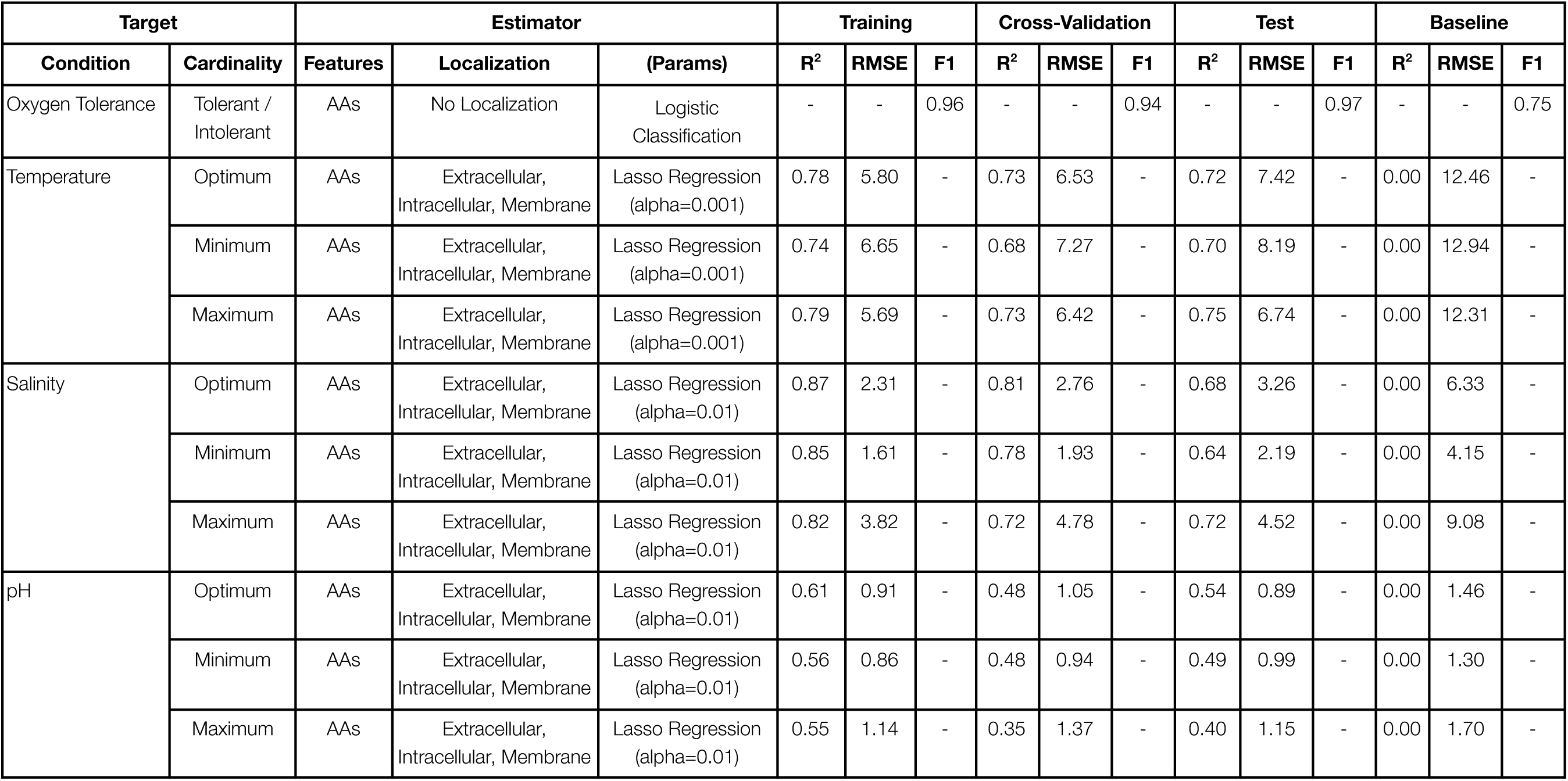
Summary of the selected models. The selected features and estimators for each model are indicated. Model performance is described for four different datasets: training, accuracy of predictions for the training data itself (*i.e.* fit of the model to its training data); cross-validation, accuracy of predictions on the examples not used to train the model during each fold of the cross-validation; test, accuracy of model on held out test data not used for model training or selection; baseline, the accuracy of a baseline prediction, which is the average of the dataset for regressions and the mode of the dataset for classifiers. Performance can be assessed using the coefficient of determination (R^2^) and root mean square error (RMSE) for continuous values and the F1 score (F1) for classifications.

We further investigated the prediction of oxygen tolerance from amino acid frequencies. Different amino acid frequencies between aerobes and anaerobes have been noted in early comparative genomics analyses^22^. Such differences could be due to phylogenetic confounders, as several large phyla are predominantly aerobes or anaerobes and differ in G+C content. Yet the model is able to discriminate between aerobes and anaerobes with a balanced accuracy of >90% from 35% to 65% G+C content, with 70% balanced accuracy outside of this range (**Supp. Fig. 3A**). The model also accurately predicts aerobes and anaerobes across prokaryotic phyla regardless of if they were composed primarily of aerobes or of anaerobes (mean 90% accuracy across phyla) (**Supp. Fig. 3A**). When the model was trained on a single phylum with equal proportions of aerobes and obligate anaerobes, *Bacteroidota* (n=419 genomes), predictions remained accurate for other phyla (F1=0.92) (**Supp. Fig. 3B**). Even two amino acids – cysteine and histidine, tryptophan, glutamate, or glutamine – can predict oxygen tolerance accurately (F1>0.90) (**Supp. Fig. 4**). Overall, this simple logistic regression model based on amino acid frequencies, which applies one set of rules across all organisms, provides a very accurate prediction (92% balanced accuracy) of oxygen tolerance generalizable to almost all bacteria and archaea.

We observed model predictions to be less accurate in some ranges of growth conditions. Optimum temperature was poorly predicted below 15°C (RMSE 14°C in cross-validation) (**Fig. 2A**). Optimum pH was systematically overpredicted below pH 5 and underpredicted above pH 9.5 (RMSE 2.0 and 1.8) (**Fig. 2A**), meaning the true pH optimum in those ranges may be more extreme than predicted. Optimum salinity was poorly predicted at 10-20% w/v NaCl for organisms that are not “salt-in” halophiles with extremely acidic proteins (RMSE 4.8% w/v NaCl) (**Fig. 2A**). At extreme growth conditions, such as temperatures ranging 60-80°C or salinity above 15%, the shift in amino acid frequencies results in a less accurate prediction of oxygen tolerance (balanced accuracy of 75% and 67%, respectively) (**Supp. Fig. 3A**).

The oxygen model contradicted reported classifications which warrant further investigation. Genera with a mix of aerobes and anaerobes accounted for 26% of instances where the oxygen tolerance model was incorrect. Specific pathways such as anoxygenic phototrophs and anaerobic sulfur oxidizing bacteria were often incorrectly predicted to be aerobes. Specific phyla such as *Campylobacterota* and *Chloroflexota* were also less accurately predicted. We hypothesize that the source of these inconsistencies may be due to inaccurate literature reports or exceptions to this model. Follow-up studies to examine the relationship between amino acid frequencies and metabolic niches will be required.

### Predictive models are robust to phylogenetic bias

Next, we evaluated the influence of phylogenetic bias on all models. A high degree of phylogenetic bias will result in highly accurate prediction for related species (*e.g.* in the same genus or family) and less accurate prediction for distant relatives (*e.g.* phylum and class). We repeated model training and evaluation using holdouts at increasing taxonomic ranks. Our results show that phylogeny had a relatively small effect on the accuracy of our sequence-based models for oxygen tolerance, temperature, salinity, and pH (**Fig. 2C**).

As a point of comparison, we show that using the same data, models strongly influenced by phylogeny fail to predict growth parameters at higher taxonomic ranks and are only accurate for close relatives (**Fig. 2C**, **Supp. Fig. 5**). These models, which strongly depend on the composition of the training dataset and the nature of phylogenetic correlation, are therefore more useful for interpolation (predictions for taxa like those in model training) than extrapolation (predictions for new, different taxa). For example, the phylogeny-based model of oxygen tolerance retained a moderate overall accuracy up to order and even class levels (**Fig. 2C**) because of the conservation of oxygen tolerance across large clades of organisms^2^ and the presence of many organisms in those clades in the dataset. Overall, these results highlight the importance of holding out test data at higher taxonomic levels to evaluate model performance^18,32^. The accuracy of our models on higher taxonomic ranks supports their use on phylogenetically novel organisms.

### Prediction using incomplete genomes

Models based solely on sequence features require a representative portion of a genome to provide accurate predictions, and can thus be applied to partial genome sequences. This is particularly meaningful for metagenome-assembled genomes, which can often be incomplete, and for genomes with non-standard genetic codes, where protein predictions may be incorrectly truncated To determine the effect of genome completeness on model performance, we randomly subsampled the proteins and contiguous fragments of a genome, independently, to 10-100% completeness. We performed this test on 20 different species spanning each condition’s range and found that our models, other than pH, retained an acceptable accuracy down to 10% genome completeness (**Fig. 2D, Supp. Fig. 6**). Specifically, we found the mean difference between 10% and 100% genome completeness to be negligible for oxygen tolerance (change in probability of 0.02), 4°C temperature, and 0.7% w/v NaCl salinity. Prediction of pH was substantially less accurate, with an error of 0.4 pH units at 10% genome completeness. Overall our models are compatible with the 50% completeness threshold used by many metagenome-assembled genome studies and the Genome Taxonomy Database^33^.

### Prediction of physicochemical conditions for all sequenced bacteria and archaea

Of the 85,205 sequenced species of bacteria and archaea available, an estimated 70% are uncultivated^33^. Uncultivated organisms were not present in our training dataset and may have unpredictable differences compared with cultivated organisms. We thus evaluated model performance on uncultivated organisms, specifically new taxonomic groups and less-complete, metagenome-assembled genomes. We first quantified the similarity of predictions for cultivated and uncultivated organisms in the same taxonomic family. We reasoned that at the family level, species might differ in cultivability while typically preferring similar growth conditions. We compared all 820 families with both cultivated and uncultivated species in the Genome Taxonomy Database, where the average family consists of 62% uncultivated species (range between 0.6-99.7%). We found high correlation (R^2^=0.8-0.9) between predicted values for cultured and uncultured microbes across all models (**Fig. 3A**), suggesting that our model can robustly predict growth conditions for uncultivated species. The high correlation for oxygen (R^2^=0.91) and relatively low correlation for pH (R^2^=0.79) agree with the expected phylogenetic conservation of those growth conditions^2,11^.

**Figure 3.**
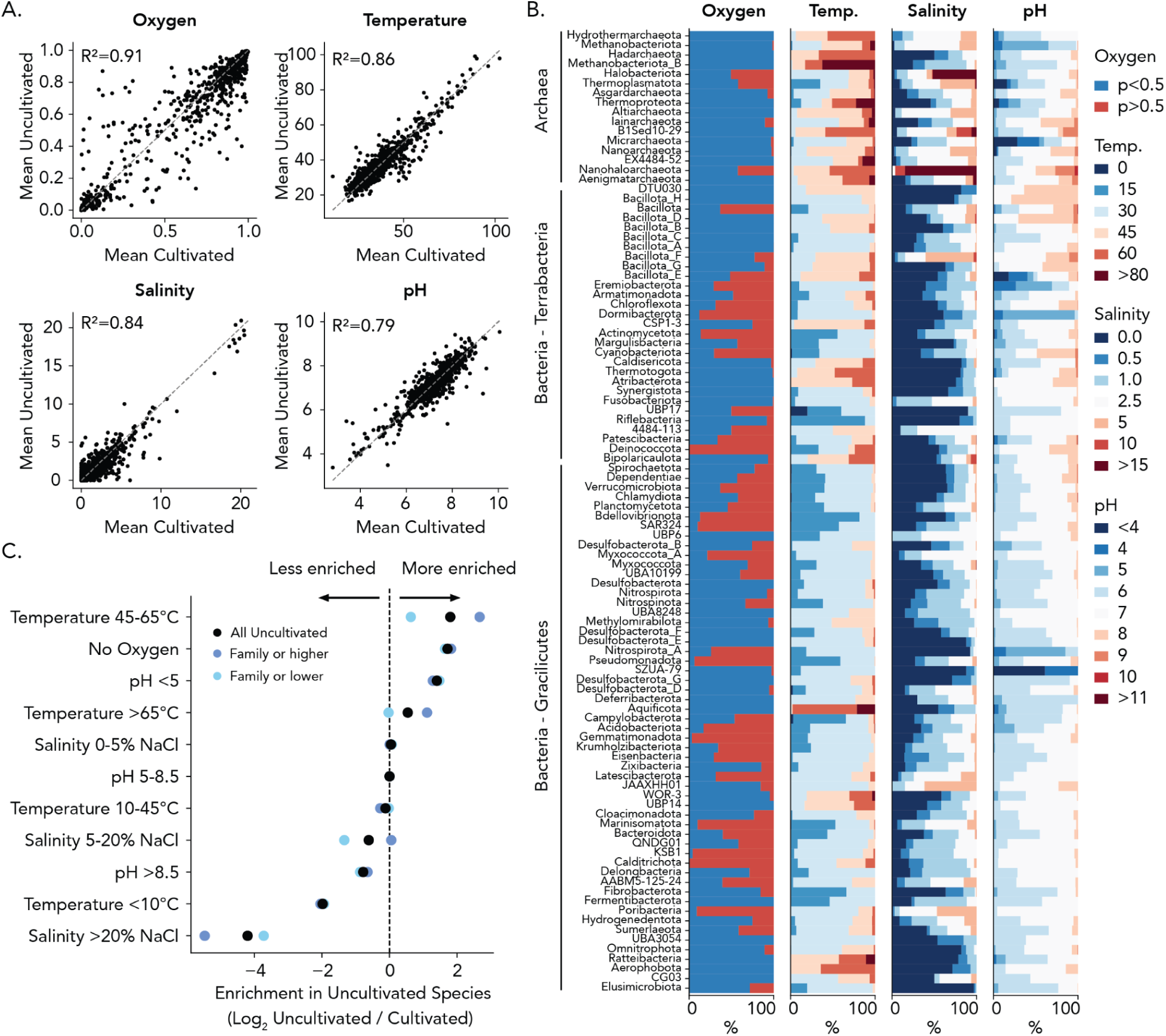
Predicted physicochemical growth conditions for 85,205 sequenced species of Bacteria and Archaea. **(A)** Evaluation of prediction bias between cultivated (x-axis) and uncultivated (y-axis) species. Correlation is shown between the average value for cultivated vs. uncultivated species in the same family, reported in coefficient of determination. Most values for oxygen tolerance are near 0 and 1. **(B)** Proportion of species predicted to grow at a given growth condition. Each plot represents one physicochemical growth condition. Each row on the y-axis represents a phylum with 10 or more sequenced species, shown in phylogenetic order starting with Archaea. The x-axis represents percent of all tested species. Each color represents a bin of values for this growth condition (e.g. “pH 5” is an optimum from pH 5-6 and “Oxygen p>0.5” are predicted aerobes). **(C)** Comparison between cultivated and uncultivated organisms predicted to grow at extreme growth conditions. For each physicochemical parameter (y-axis), the fold change (log2 scale) between cultivated and uncultivated species is shown. Positive values indicate the proportion of uncultivated species with those conditions is greater than the proportion of cultivated species. Color indicates results for different groups of uncultivated species: black - all uncultivated species; light blue - uncultivated species in families with at least one cultivated representative; dark blue - uncultivated species in families with no cultivated relatives (i.e. uncultivated above the family level).

We found several examples of taxonomically distinct lineages suggesting our results are in agreement with previously published observations. The oxygen tolerance model correctly predicted anaerobic classes of *Actinobacteria* and the class *Vampirovibrionia* (formerly *Melainabacteria*) in the phylum *Cyanobacteriota* to be anaerobic, unlike its aerobic relatives (**Supp. Fig. 7**)^34,35^. Within the *Chloroflexota* class *Dehalococcoidia*, the model correctly identified the order *Dehalococcoidales* to be entirely anaerobic and orders *Tepidiformales*/OLB14 and SAR202/UBA1151 as containing aerobes, in agreement with previous reports^36–38^. Elevated optimal temperature was correctly predicted for phyla *Hadarchaeota* and *Calescibacterota* (**Fig. 3B**)^39,40^. Elevated salinity was correctly predicted for the phylums *Nanohaloarchaeota* (**Fig. 3B**)^41^ and *Salsurabacteriota*/T1Sed10-126 (**Supp. Data 4**), which additionally has a predicted optimum pH in line with its reported habitat in soda lakes^42,43^. In addition, a large proportion of species in the phyla *Eremiobacteriota*, SZUA-79 (monotypic order *Acidulodesulfobacterales*), and *Micrarchaeota* were predicted to grow at low pH (**Fig. 3B**), consistent with previous assessments^10,44,45^. These examples support our hypothesis that sequence feature-based models can learn from cultivated microbes and be generalized to predict physicochemical requirements for uncultivated groups of life.

We used the models to compare overall growth conditions between uncultivated microorganisms and those cultivated thus far (**Fig. 3C**, **Supp. Fig. 8**). Perhaps unsurprisingly, we found that uncultivated organisms are more likely to be thermophiles, anaerobes, and acidophiles. Specifically, uncultivated organisms were 3.5-fold more likely to grow at temperatures between 45-60°C, 3.3-fold more likely to grow in anoxic conditions, and 2.6-fold more likely to grow at pH below 5. Anaerobes accounted for 54% of uncultivated species but only 16% of cultivated species (**Supp. Data 4**). Microorganisms in families lacking any cultivated members were 4.1-fold more likely to be thermophiles relative to uncultivated microorganisms with cultivated relatives (**Fig. 3C**). For example, 18% of species in uncultivated families were predicted to grow optimally between 45-60°C, compared to only 4% of species in cultivated families (**Supp. Data 4**). While the census of microbial life from metagenomes is incomplete and biased, these data support the notion that successful cultivation of diverse lineages may require development of accessible laboratory methods for manipulation at a broad range of growth conditions.

### Prediction of physicochemical conditions for metagenomes across habitats

A key consideration for cultivating microorganisms from a given habitat is the design of a laboratory environment to support their growth. Often, growth conditions are chosen that closely resemble the microorganism’s habitat. In order to assess the heterogeneity of growth conditions between and within organisms sharing the same habitat, we predicted optimum growth conditions for a publicly available dataset of 52,515 metagenome–assembled genomes from various habitats (**Supp. Data 5**)^46^. We compared all 3,349 metagenomes that contain five or more genomes, of which 37% are from vertebrate hosts, 21% are from marine environments (includes waters, sediments, and hydrothermal vents), 12% are from freshwater, and 5% are from soil. We found the average data across metagenomes generally aligns with expected habitats trends (**Fig. 4A**). For example, vertebrate-related metagenomes, predominantly from gut samples, were largely anaerobic and displayed limited variation in pH (median 7.1) and temperature (median 36°C) (**Fig. 4A**). As expected, the median of the average optimum temperatures of thermal springs metagenomes and thermal marine metagenomes was 69°C and 62°C, respectively. Similarly, metagenomes with a mean predicted optimum salinity above 6% w/v NaCl, 81% were from non-marine saline and alkaline or engineered environments. Metagenomes with mean predicted pH of 5.0 or less were from thermal springs, ferrous or sulfidic biofilms, peatlands, bogs, or “freshwater.” Therefore, our models can extend laboratory based measurements of microorganisms to estimate environmental conditions.

**Figure 4.**
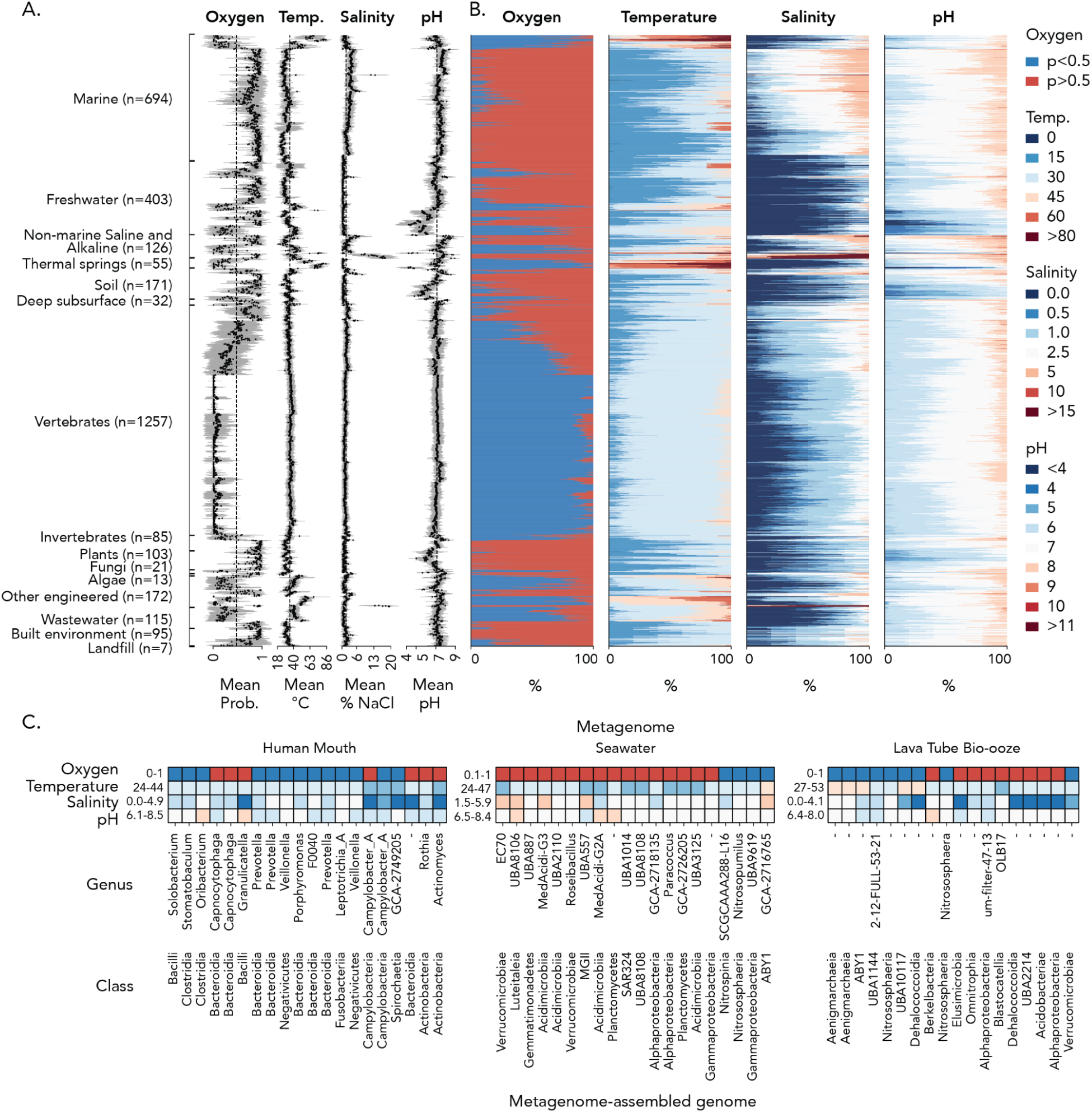
Predicted physicochemical growth conditions for metagenome-assembled genomes from 3,349 metagenomes. **(A)** Predicted values for each growth condition in each metagenome, with the mean shown as black dot and ±1 standard deviation shown in gray. Metagenomes (y-axis) were sorted by clustering within each biome (label). The number of metagenomes in each biome is shown in parentheses. The vertical line is the average values across all metagenomes in this dataset. **(B)** Distributions of predicted growth conditions among genomes in each metagenome. Each color represents a bin of values for this growth condition (e.g. “pH 5” is an optimum from pH 5-6 and “Oxygen p>0.5” are predicted aerobes). **(C)** Examples of how combinations of predicted growth conditions can vary for specific organisms within three metagenomes with 20 metagenome-assembled genomes each. Colors correspond to those used in (B) and text indicates genus and class as assigned in the GEM dataset (some taxonomic levels are unassigned). Minimum and maximum values for each condition are listed next to each metagenome.

Yet within a metagenome, predictions for individual genomes can vary from the average. For example, metagenomes with mostly obligate anaerobes, like those from vertebrate guts, often contained aerobes and metagenomes with mostly aerobes, like those from marine environments, sometimes contained obligate anaerobes (**Fig. 4B**). Across all metagenomes, the difference between the minimum and maximum predicted optima for organisms in a metagenome had a median value of 16°C, 3.2% NaCl, and 1.6 pH units. Possible explanations for this heterogeneity, other than prediction error, could include a combination of heterogeneous conditions at the microscopic scale, environmental fluctuations, transient dispersal, dormant or non-growing cells, and relic DNA. We anticipate genome-based predictions of growth conditions could help microbial ecologists understand in more detail ecological processes contributing to heterogeneity of optimal conditions, such as migration and environmental fluctuation. Because predictions are made on the genome level, researchers can identify microorganisms in communities with different combinations of growth conditions (**Fig. 4C**). Such predictions may suggest the use of several different growth conditions to isolate more diverse members of the community, or in select cases even help isolate a specific microorganism by suggesting conditions supporting their distinct growth.

## Discussion

This work provides a set of models for predicting physicochemical growth conditions of microorganisms based solely on genome features. A key aspect of these models is their robustness to phylogenetic bias, which allows accurate prediction of multiple taxonomic levels using a microbe’s genome sequence. We also demonstrate robust prediction using partial genome sequences that meet the standards of metagenomic analysis (>50% complete). This data allows an unprecedented examination of predicted physicochemical conditions across presently sampled bacteria and archaea (**Supp. Data 4**). This is of critical importance for ongoing efforts to cultivate diverse microorganisms^47,48^. We envision that the physicochemical models provided here will contribute to cultivation of uncultured organisms, whose characterization in turn will improve the models. The models could also be used to understand points in the tree of life where organisms transitioned from one growth condition to another, such as anaerobe-aerobe transitions ^16^.

The ability to predict if a microbe is an aerobe or anaerobe based solely on the frequencies of two or more amino acids is an unexpected finding that warrants further study. Differences in the frequency of oxygen-sensitive amino acids between aerobes and anaerobes were initially attributed to reactivity with oxygen, but this hypothesis has been disputed^22,23,49^. We suspect the correlation between oxygen tolerance and amino acids across phyla (**Supp. Fig. 9A**) is largely attributable to the cost of biosynthesis. Tryptophan and histidine have higher energetic costs than similar amino acids^50^ (**Supp. Fig. 9B**), and would be relatively more costly to synthesize in anoxic environments where less energy is available. Cysteine is significantly more costly in oxic environments because of the extra energy needed for cells to use the oxidized forms of cysteine and sulfur^51^. Prediction of facultative anaerobes must be identified from genes associated with anaerobic respiration or fermentation. Facultative anaerobes and aerobes have similar amino acid usage, indicated by probability of oxygen tolerance (**Supp. Fig. 9C**) and similar overall gene content^12^. A recent effort was unable to accurately predict obligate aerobes, facultative anaerobes, and obligate anaerobes using amino acid frequencies (balanced accuracy 55%)^21^. The similarity between facultative anaerobes and aerobes is evidence of the strong selective pressure that oxic conditions exert on amino acid usage. Studies of organisms where predicted oxygen tolerance differs from these identified amino-acid usage patterns (*e.g.* obligate anaerobes predicted to be oxygen tolerant) would be of particular interest to better understand selective pressures on specific metabolisms or taxa.

This work represents a significant improvement over existing resources for predicting physicochemical growth conditions (**Table 4**). Our models predict all conditions at once and include optimum, minimum, and maximum. By predicting continuous values, they provide more precision than models predicting categories (*e.g.* “thermophile”). As sequence-based models, they compute more quickly than models involving gene annotations. The composition of all training datasets are biased entirely by cultivated organisms, but some published models have other biases that may impact performance: the best performing oxygen model did not include facultative anaerobes in model training and evaluation^16^, and the best performing optimum pH model was trained and evaluated specifically on soil and freshwater bacteria^11^. A key concern with many models with apparent high performance is that they used high-dimensional features prone to overfitting on training data (*e.g.* hundred to thousands of genes or k-mers) yet, except for one such model^16^, did not use out-of-clade test data to make sure predictions remained equally accurate for organisms in unrelated taxa, which is unlikely. We expect that modeling efforts will continue to improve and that the simple, validated sequence-based models presented here will provide a useful foundation for that research.

**Table 4.** Comparison of published models predicting oxygen, temperature, salinity, and pH. The following parameters are shown for each study: prediction type; cardinality, the number of things being predicted; features, the variables used for the prediction (k refers to k-mer size); training data composition, what organisms were used (bias); feature size, the number of features; training data size, the number of examples learned; balanced taxonomically, whether the publication reports intentionally reducing taxonomic bias to a significant degree (not simply de-replicating to 1 genome per species); tested, if the accuracy is measured using examples not used in model training and selection; test out-of-clade, if taxonomic groups were intentionally held out for testing; test accuracy, score to test performance; accuracy metric, metric used to score performance - accuracy is fraction correct assignments, balanced accuracy is mean of recall for all predicted categories, F1 is harmonic mean of the precision and recall, R^2^ is coefficient of determination; study, publication.

| Condition | Prediction Type | Cardinality | Features | Training data composition | Feature size | Training data size | Balanced | Tested | Tested out-of-clade | Test accuracy | Accuracy metric | Study |
| --- | --- | --- | --- | --- | --- | --- | --- | --- | --- | --- | --- | --- |
| Oxygen | Category | Aerobe (Any) or Obligate Anaerobe | Protein (k=1) | Isolates | 20 | 3646 | Yes | Yes | Yes (Family) | 0.91 | Balanced accuracy | This work |
| Oxygen | Category | Aerobe, Facultative Anaerobe, or Obligate Anaerobe | Protein (k=3) | Isolates | 8000 | 3350 | No | Yes | No | 0.7 | Balanced accuracy | Flamholz<sup>21</sup> |
| Oxygen | Category | Aerobe, Facultative Anaerobe, or Obligate Anaerobe | DNA (k=5) | Isolates | 512 | 3350 | No | Yes | No | 0.67 | Balanced accuracy | Flamholz<sup>21</sup> |
| Oxygen | Category | Aerobe or Anaerobe | Genes | Isolates, aerobes and anaerobes | 3507 | ~3200 | No | Yes | Yes (Family) | 0.97 | Accuracy | Davin<sup>16</sup> |
| Oxygen | Category | Aerobe or Anaerobe | Sequence alignment | Isolates, aerobes and anaerobes | 99 | 333 | Yes | Yes | No | 0.84 | Accuracy | Davin<sup>16</sup> |
| Oxygen | Category | Aerobe or Anaerobe | Genes | Isolates | 542 | 5194 | No | Yes | No | 0.91 | Balanced accuracy | Reimer<sup>1</sup> |
| Oxygen | Category | Aerobe, Facultative Anaerobe, or Obligate Anaerobe | Genes | Isolates | 325 | 272 | Yes | Yes | No | 0.8 | Accuracy | Jabłońska<sup>12</sup> |
| Oxygen | Category | Aerobe, Facultative Anaerobe, or Obligate Anaerobe | Genes | Isolates | 44498 | 549 | No | Yes | No | 0.82 | Accuracy | Edirisinghe<sup>14</sup> |
| Oxygen | Category | Aerobe, Facultative Anaerobe, or Obligate Anaerobe | Genes | Isolates | 6554 | 2231 | No | Yes | No | 0.86 | F1 | Weber Zendrera<sup>15</sup> |
| Oxygen | Category | Aerobe or Anaerobe | Genes | Isolates, mostly pathogens | 14831 | 234 | No | Yes | No | ~0.97 | Accuracy | Weimann<sup>13</sup> |
| pH | Value | Opt., Min., and Max. | Protein (k=1) | Isolates | 60 | 756 | Yes | Yes | Yes (Family) | 0.54 | R<sup>2</sup> | This work |
| pH | Value | Habitat Preference | Genes | Uncultivated soil and aquatic genomes | >335 | 580 | No | Yes | No | 0.55 | R<sup>2</sup> | Ramonedá<sup>11</sup> |
| pH | Category | Acidophile or not | Genes | Isolates | 542 | 2710 | No | Yes | No | 0.67 | Balanced accuracy | Reimer<sup>1</sup> |
| Salinity | Value | Opt., Min., and Max. | Protein (k=1) | Isolates | 60 | 593 | Yes | Yes | Yes (Family) | 0.68 | R<sup>2</sup> | This work |
| Salinity | Category | Halophile or not | Genes | Isolates | 542 | 1162 | No | Yes | No | 0.98 | Balanced accuracy | Reimer<sup>1</sup> |
| Temperature | Value | Opt., Min., and Max. | Protein (k=1) | Isolates | 60 | 1722 | Yes | Yes | Yes (Family) | 0.72 | R<sup>2</sup> | This work |
| Temperature | Value | Optimum | tRNA (k=1) | Isolates | n.r. | 783 | No | Yes | Yes (Clusters) | 0.59 | R<sup>2</sup> | Cimen<sup>32</sup> |
| Temperature | Value | Optimum | DNA (k=2-8) | Isolates | ~44000 | 1815 | No | Yes | No | 0.88 | R<sup>2</sup> | Wang<sup>66</sup> |
| Temperature | Value | Optimum | Protein (k=2) | Isolates | 400 | 5762 | No | Yes | No | 0.9 | R<sup>2</sup> | Li<sup>67</sup> |
| Temperature | Value | Optimum | Protein (k=1) | See Zeldovich | n.r. | n.r. | n.r. | n.r. | Yes (Metagenomes) | 0.87 | R<sup>2</sup> | Kurokawa<sup>26</sup> |
| Temperature | Value | Optimum | Protein, DNA, and RNA (k=1) | Isolates | 29 | 2719 | No | Yes | No | 0.84 | R<sup>2</sup> | Sauer<sup>24</sup> |
| Temperature | Value | Optimum | Protein (k=1) | Isolates | 20 | 84 | No | Yes | No | 8-9 | RMSE | Zeldovich<sup>25</sup> |
| Temperature | Value | Optimum | Genes | Isolates | 6554 | 782 | No | Yes | No | 0.77 | R<sup>2</sup> | Zendrera<sup>15</sup> |
| Temperature | Category | Thermophile or not | Genes | Isolates | 542 | 8406 | No | Yes | No | 0.79 | Balanced accuracy | Reimer<sup>1</sup> |

One issue with all available predictions of optima is that they are likely not precise enough to help improve laboratory work with cultured microorganisms. The exception would be situations where the physicochemical conditions used for isolation are not reasonably close to optimum conditions for growth. When using models, researchers should be aware of the average error as well as ranges with higher error (for example, below 25°C and at extreme pH values with our models) and that unusual organisms could have much greater error. Predictions of minimum, maximum and optimum can disagree when predicted independently (*e.g.* maximum equal or less than minimum), as they do for our salinity predictions for 5% of the GTDB dataset. A key source of error is the imprecision of the training data, which is biased towards particular intervals (*e.g.* 25°C, 30°C, 37°C, 42°C). Collection of higher-granularity phenotypic data will help improve models and is especially important to measure for novel taxa that are more likely to have unusual features^52^.

The use of sequence features might be improved by incorporating structural information. For example, protein isoelectric point would be most accurately calculated using residues on the protein surface but is currently calculated using all residues in the protein sequence. Structural predictions could also enable the calculation of new, more informative features, such as interactions between residues influencing temperature stability^53^. Such structural information could be needed for accurate prediction of low optimum temperatures, where adaptations to increase protein flexibility are varied^54^. We expect improvements in structural datasets to support more accurate computational models for microbiology.

Future efforts can build on sequenced-based predictions to make more accurate gene-based models. A limitation of sequenced-based models is that, relative to changes in genes, changes in amino acid composition are slow and sometimes subtle. For example, gene gain and loss enable transitions between freshwater and seawater that take millions years to become evident in amino acid composition^27^. Similarly, an organism that acquires oxygen-sensitive genes and becomes oxygen intolerant would initially continue to have the amino acid composition of an oxygen tolerant organism. Small shifts in optimum pH can have significant impacts in the solubility and protonation of specific chemicals, and therefore on the fitness effect of specific genes^55–57^, yet result in only subtle change to amino acid composition. Despite these potential caveats, gene-based models do not currently provide a better alternative because in most cases the genes underlying growth conditions in diverse organisms are unknown. A promising approach would be using the sequence-based predictions of growth conditions now available for all sequenced cultivated and uncultivated organisms to find phylogenetically robust correlations between conditions and genes. Such genes would be an appropriate set of features for building accurate gene-based models.

## Conclusions

Microbial adaptation to physicochemical growth conditions of oxygen concentration, temperature, pH, and salinity leave signatures in the DNA and protein sequences of genomes that can be used to predict optimum conditions. In predicting these conditions, an important confounder is the phylogenetic bias of growth conditions among related organisms and which organisms have conditions measured to begin with. The models and approach presented here provide a foundation for future work predicting physicochemical conditions with more precision. Presently, these models provide exciting new information about the physicochemical conditions preferred by uncultivated organisms, possibly aiding in the cultivation of new lineages of life.

## Methods

### Data sources and preprocessing

Publicly available growth phenotypes were obtained from BacDive accessed by API in June, 2023^1^. Data on temperature, pH, salinity, and oxygen tolerance were coded as follows: for continuous variables, reported optimum was used, and minimum and maximum were defined as the least and greatest values where growth was described as “positive” and, for salinity only, “inconsistent.” If an optimum was reported in a range, which was common for pH, we recorded optimum as the average of that range. Oxygen tolerance was assigned as tolerant for microbes described as ‘obligate aerobe’, ‘aerobe’, ‘facultative anaerobe’, or ‘facultative aerobe’ and intolerant for microbes described as ‘obligate anaerobes’ or ‘anaerobes’.

Genome accessions were used to download genomes from NCBI GenBank^58^. Additional genomes for analyses were downloaded from the Joint Genome Institute Genomic Catalog of Earth’s Microbiomes (GEM) project^46^ and from the Genome Taxonomy Database (GTDB) release r214, which also supplied the taxonomy used here to describe organisms and to identify nearest taxonomic neighbors^33^. When not available, predicted coding domain sequences (CDS) were identified from genomes based on open reading frames using the standard genetic code using Prodigal v.2.6.3 with default settings^59^.

Preprocessing used an automated curation to remove lower quality phenotypic data. For example, a study that measured temperature in 1°C increments from 20-40°C is of higher quality than a study that only measured 30°C, 37°C, and 40°C. Preprocessing sought to deplete the dataset of lower quality phenotypic data like the latter. For commonly measured optima, defined as optima that accounted for more than 1% of values (*e.g.* strains with optimum pH 7.0 accounted for 34% of pH values), data were removed if the minimum or maximum equaled the optimum; if the difference between the minimum and maximum was less than 1.5 pH units, 10°C, or 1.5% NaCl (unless salinity was <0.5% NaCl); or if fewer than 4 total points were reported. More extreme values – pH <4 or >9, salinity >15% NaCl, and temperature <19°C or >44°C – were not removed as we reasoned these are more likely to be accurately measured. Data was discarded if any of the optimum, minimum, or maximum were not reported. Due to the unique potential for data entry errors with salinity, we discarded organisms from the dataset if they were haloarchaea with a reported optima under 3.7% or if they were reported to have an extreme salinity (above 14% NaCl) and were obviously incorrect. We expect that inconsistencies in definitions and testing of oxygen requirements, such as not testing anaerobes for facultative aerobic growth, mean that oxygen requirement data also contains errors, but these were not addressed.

### Measurement of genomes

Properties of proteins and DNA sequences for each genome (**Table 1**) were measured using custom code. Measurements were made on individual predicted mature proteins, which are protein sequences where the N-terminal methionine and any signal peptide are removed. Protein measurements included amino acid frequencies, predicted isoelectric point^60^, Grand Average of Hydropathy (GRAVY)^61^, and average carbon oxidation state (Zc) and stoichiometric hydration state (nH2O)^62^. Measurements were then aggregated across proteins in the genome. Measurements made on DNA are more limited: G+C content, the frequency of purine-pyrimidine transitions, and the estimated coding density. Genomes were discarded if their coding density was below 60%. To obtain genomes of varying completeness, proteins were randomly sampled and each DNA sequence (*e.g.* contig) had a random contiguous portion sampled according to the desired level of completeness.

The signal peptide prediction model used above was developed to identify exported proteins very quickly and off-license to support fast use of the models without use limitations. A hidden Markov model was constructed as described in published methods reported to be accurate at discriminating the presence or absence of signal peptides^63–65^. The model has states for positions in the start, N-terminal, hydrophobic, and cut-site regions of SPase I signal peptides, and the mature protein sequence and was fit using a published dataset^64^. The model runs very quickly and has high specificity and sensitivity. Only about 10% of proteins are exported, however, so a threshold was chosen to only allow 20% of proteins in a genome to be false positives, which captures 60% of exported proteins in the genome (**Supp. Fig. 10**). Proteins are assigned as a localization as follows: “membrane” is the protein’s GRAVY is greater than 0.5 the proteome’s average GRAVY, otherwise “intracellular soluble” if the protein lacks a signal peptide and “extracellular soluble” if the protein has a signal peptide.

### Model dataset construction

To reduce the imbalances in the dataset towards particular groups of groups, organisms from overrepresented groups were removed. First, the frequency of taxa in the dataset was compared to the expected frequency of taxa at each taxonomic level. The fold enrichment in the dataset is equal to the ratio between observed and expected frequencies for each taxonomic level multiplied together. Then, a random portion of taxa is selected, with the probability of being selected inversely proportional to their fold enrichment. The result is a smaller dataset more similar to the expected composition. Here we used the composition of representative species in the Genome Taxonomy Database as the expected composition and the portion of data removed was 50% for oxygen data and 33% for every other condition, which had more extensive curation. Extreme values were retained as in data preprocessing.

The partitioning of data into training and test datasets was taxonomically aware. Random families totaling 20% of the curated, balanced dataset were held out to make a test dataset, and the remaining 80% of the dataset was for model training and cross-validation. To ensure the test dataset had a wide range of data, partitioning at the family level was done independently for extremely low values (<3rd percentile), extremely high values (>97th percentile), and the remaining values (3rd-97th percentiles). This introduces some data leakage in the sense that the training and test set can include members from the same families, but the members in the test set have extreme traits, whereas the members in the training set do not. In cross-validation, random families consisting of approximately 20% of the training dataset were held out in each fold.

### Model evaluation and selection

Different combinations of features and estimators (untrained models) were tested through cross-validation to select the features and estimator to use to predict each growth condition (**Supp. Data 3**). Nine sets of features tested included three types of variables – amino acid frequencies, pI distributions, other more-derived features, or all features – and three different compartmentalizations – no compartmentalization, by compartment (*e.g.* intracellular, extracellular, and membrane proteins individually), and differences between extracellular and intracellular proteins. For all models, standard scaling was applied to the features. Feature selection of either the 20 most correlated features (measured with the F statistic or mutual information statistic) or the minimum number of features were used for most models. Regression estimators tested were standard, ridge, and lasso (no feature selection) linear regressions, with varying hyperparameters, and multi-layer perceptron regressor with two layers and max features per layer of half the available features (no feature selection). Classifier estimators tested were Gaussian naive Bayes, logistic regression, and support vector machine (no feature selection) with a linear kernel, with varying hyperparameters. Each fold of the cross-validation involved its own scaling, feature selection, and estimator fitting.

Regression model performance was assessed using the coefficient of determination (R^2^) and root mean square error (RMSE). Classification model performance was assessed on true positives, false negatives, etc., using specificity, sensitivity, the harmonic mean of those values (F1), and true and false positive rates. For model selection, the above combinations of models and features were compared using R^2^ or F1 scores on the cross-validation data (**Supp. Fig. 1**). Among the highest scoring estimators and sets of features, we selected an estimator and features based on simplicity and perceived closeness to the biological phenomena. For example, the oxygen tolerance models were equally accurate when trained on all features or on only amino acids, so we chose to use only amino acids. Feature importances in the selected models were computed with the permutation importance (**Supp. Fig. 11**). After model selection, models were retrained as before but on all training data at once (no cross-validation), then predictions were made on the final test dataset.

### Model use

A software package is available for users to predict growth conditions for genomes (<u>github.com/cultivarium/GenomeSPOT</u>). The final models were trained on all curated, balanced data from both the training and test datasets. The software reports predicted optima, max, and min and oxygen tolerance. The runtime is approximately 5-10 seconds per genome per CPU.

## Author Contributions

T.B., A.C.C., H.H.L., and N.O. conceived of and designed the project. T.B., A.C.C., and M.M. designed model training and evaluation. T.B. produced all code, data, and results. T.B., A.C.C., M.M., P.C., H.H.L., and N.O. analyzed results and wrote the manuscript. All authors have reviewed the manuscript and approved of its release.

## Acknowledgements

We thank the entire Cultivarium team for their support in this work. We are especially grateful to Elise Ledieu-Dherbécourt for advice and support on the project. We recognize Paul Carini for the insight that the influence of oxygen amino acid composition may be explained by amino acid biosynthesis costs. We extend special thanks to James Knight for code support and review and Julia Leung and Stephanie L. Brumwell for their feedback on the manuscript. Cultivarium acknowledges support from Schmidt Futures as a Convergent Research Focused Research Organization (FRO).

## Supplementary Information

**Supplementary Figure 1.** Model selection experiment results for each condition.

**Supplementary Figure 2.** Performance for minima, optima, and maxima

**Supplementary Figure 3.** Additional evaluation of the oxygen model.

**Supplementary Figure 4.** Evaluation of oxygen models using only two features.

**Supplementary Figure 5.** Holdout experiment scatter plots

**Supplementary Figure 6.** Accuracy vs. genome completeness for individual genomes.

**Supplementary Figure 7.** Examples of oxygen classifications for uncultivated lineages.

**Supplementary Figure 8.** Trait distribution by cultivation status.

**Supplementary Figure 9.** Understanding the oxygen model.

**Supplementary Figure 10.** Construction of a hidden Markov model to predict signal peptides.

**Supplementary Figure 11.** Feature importances in selected models for top 20 features.

**Supplementary Data 1.** Predictions and reported phenotypes for all BacDive genomes. Columns titled “reported” refer to phenotypes reported on BacDive. Columns titled “partition” indicate if a genome was used in the training or test set. Units correspond to those used in the manuscript: oxygen - the probability of being oxygen tolerant; temperature - degrees Celsius; pH - units; salinity - % w/v NaCl. GTDB taxonomy is reported. (supplementary_data_1.tsv)

**Supplementary Data 2.** The number of genomes used in the test, train, or cross-validations (cv) holdout sets for each physicochemical condition. Data is provided at the family level, which was used for model selection. (supplementary_data_2.tsv)

**Supplementary Data 3.** Performance of all features and estimators tested during model selection. (supplementary_data_3.tsv).

**Supplementary Data 4.** Predictions for all GTDB representative species and GTDB-provided taxonomy and NCBI genome category (isolate if “none”). (supplementary_data_4.tsv) **Supplementary Data 5.** Predictions for all GEM MAGs and GEM-provided taxonomy and metagenome metadata. (supplementary_data_5.tsv)

