## Supplementary Figures for "Predicting microbial growth conditions from amino acid composition"

Supplementary Figures 2
- Supplementary Figure 1. Model selection experiment results for each condition. 2
- Supplementary Figure 2. Performance for minima, optima, and maxima 3
- Supplementary Figure 3. Additional evaluation of the oxygen model. 4
- Supplementary Figure 4. Evaluation of oxygen models using only two features. 5
- Supplementary Figure 5. Holdout experiment scatter plots. 6
- Supplementary Figure 6. Accuracy vs. genome completeness for individual genomes. 7
- Supplementary Figure 7. Examples of oxygen classifications for uncultivated lineages. 8
- Supplementary Figure 8. Trait distribution by cultivation status. 9
- Supplementary Figure 9. Understanding the oxygen model. 10
- Supplementary Figure 10. Construction of a hidden Markov model to predict signal peptides. 11
- Supplementary Figure 11. Feature importances in selected models for top 20 features. 12

### Supplementary Figures

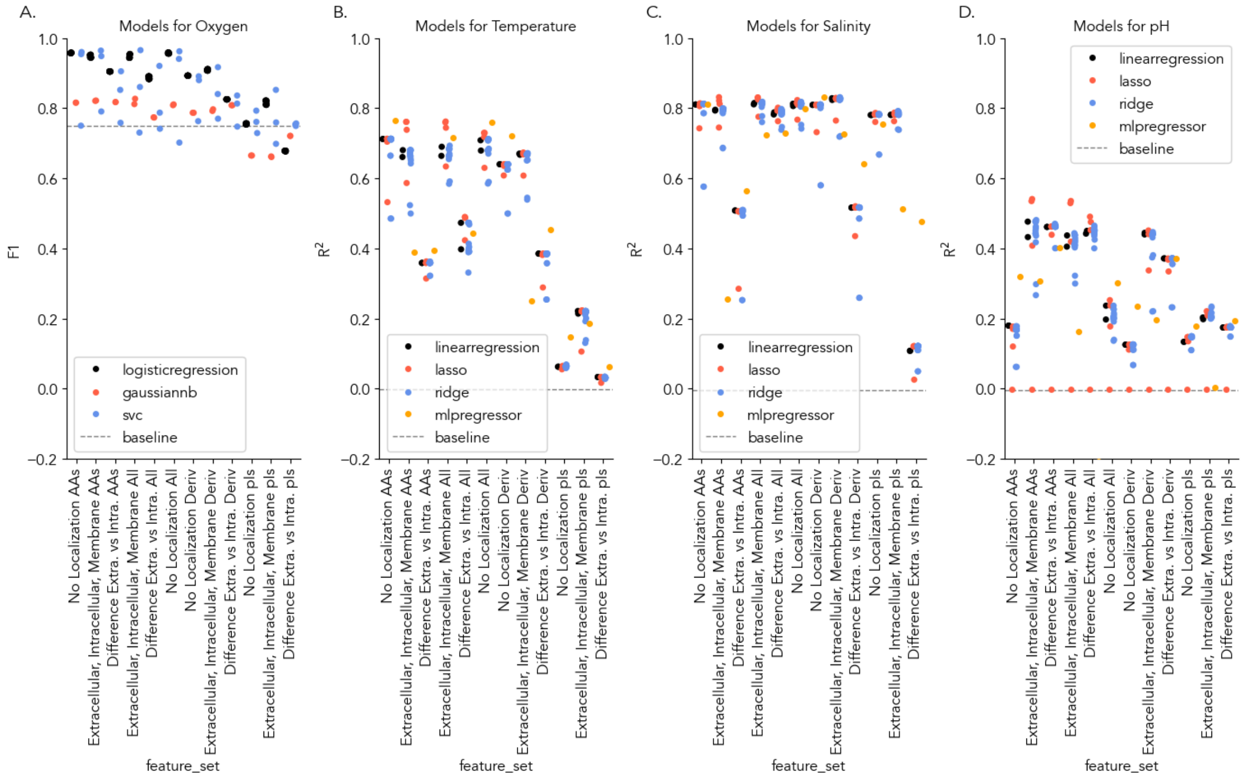


**Supplementary Figure 1. Model selection experiment results for each condition.**

(A) Oxygen tolerance, (B) optimum temperature, (C) optimum salinity, and (D) optimum pH. Each dot is an estimator trained on one set of features. Dots are grouped by type of estimator and type of features. Where multiple dots are present, the hyperparameters of the estimator have varied. A subset of these results are presented in Figure 2B. Dashed line indicates baseline values.

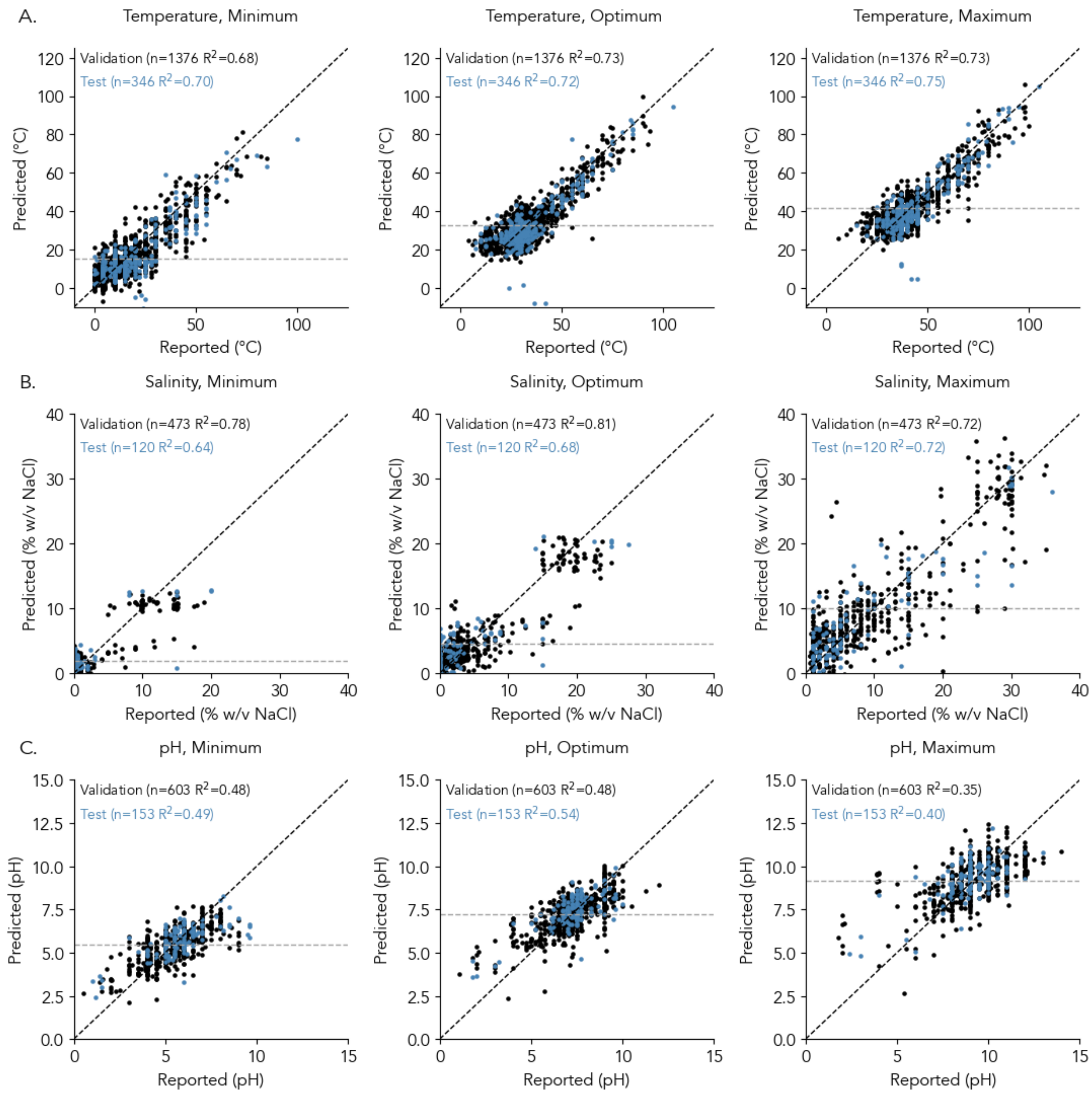


**Supplementary Figure 2. Performance for minima, optima, and maxima**

Analysis in Figure 2A expanded to show model performance for minima and maxima along with optima.

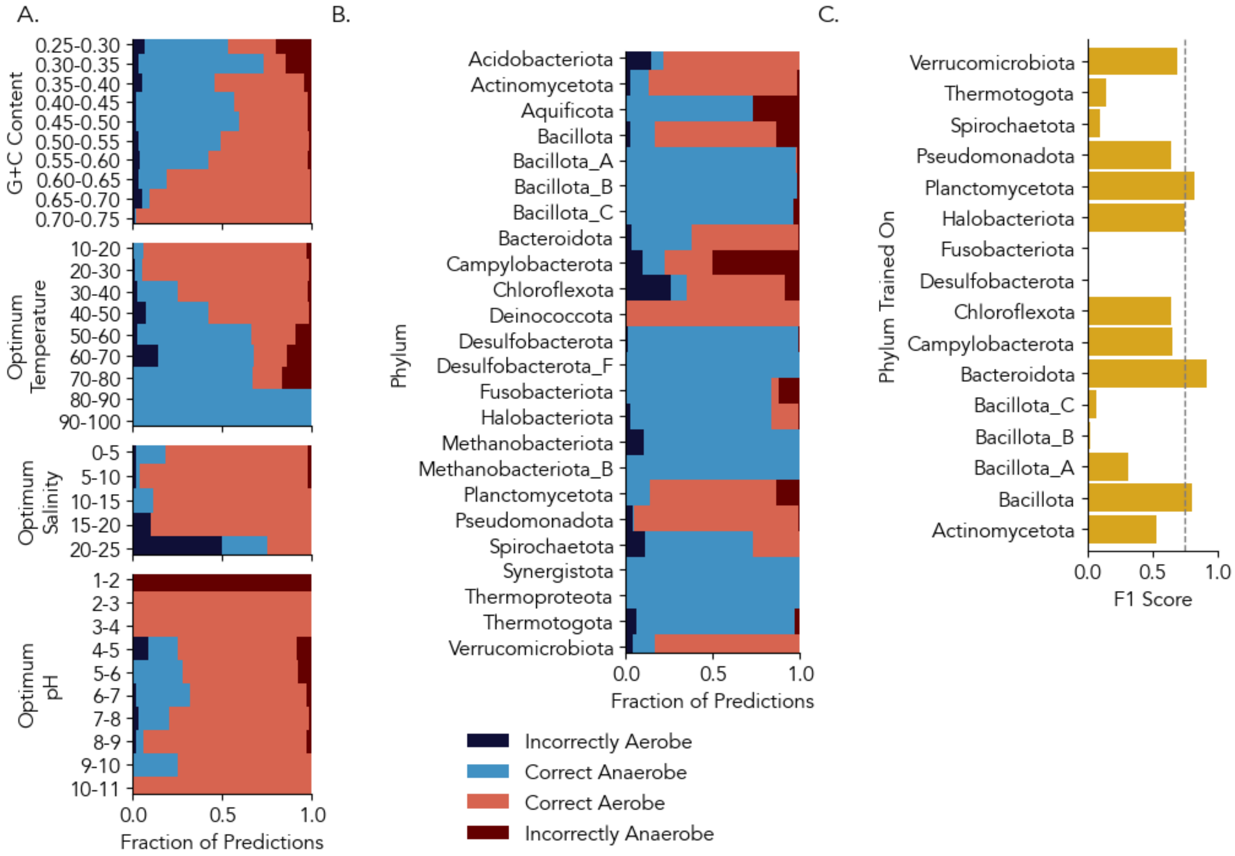


**Supplementary Figure 3. Additional evaluation of the oxygen model.**

(A-B) Each stacked barplot shows the proportion of cross-validation predictions of which oxygen tolerant microbes were correctly identified (light red) or misclassified as intolerant (dark red) and of which oxygen intolerant microbes were correctly identified (light blue) or misclassified as tolerant (dark blue). Barplots group organisms according to (1) genomic G+C content, (2) reported optimum temperature, (3) reported optimum salinity, and (4) phyla with 10 or more species in the training dataset. (C) The accuracy of the prediction on a validation dataset composed of all other phyla, when the model is only trained on species of the indicated phylum. Dashed line is the baseline for the entire training dataset, which is rarely exceeded for most phyla.

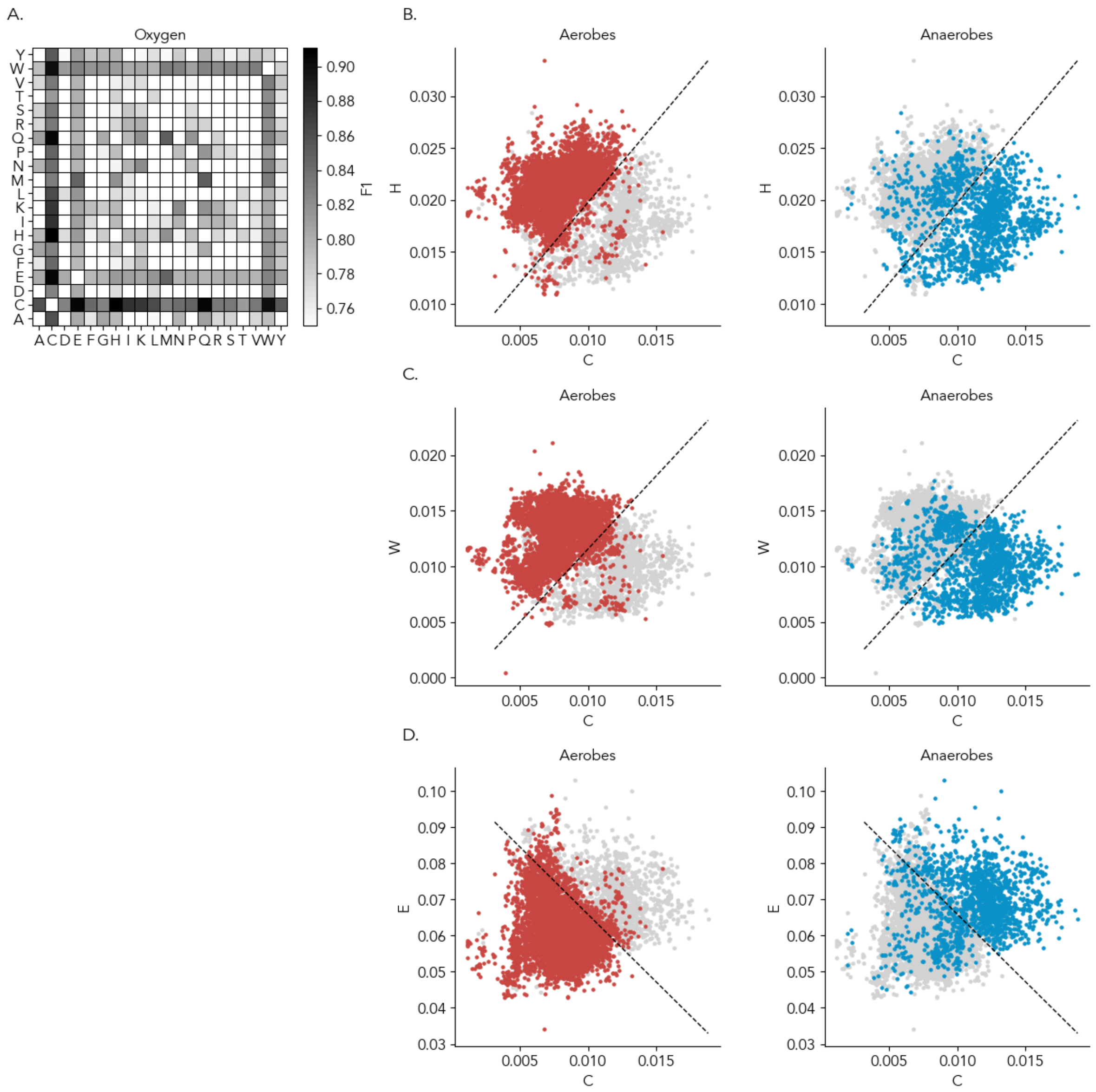


**Supplementary Figure 4. Evaluation of oxygen models using only two features.**

(A) A heatmap displaying the F1 score (colorscale) in cross-validation when a model is trained using a pair of amino acids (rows and columns). The minimum value is the baseline for the training. (B-D) Scatterplots showing each species according to cysteine frequency (x-axis) and histidine, tryptophan, or glutamate frequency (y-axis). Left plots show species described in the literature as aerobes as red dots in the foreground and right plots show species described in the literature as anaerobes as blue dots in the foreground. A natural separation between aerobes and anaerobes can be observed. The dashed line indicates the decision boundary for a logistic regression trained on the dataset, which automatically separates the two groups. Decision boundaries can be reproduced with (eq. 1) H = 1.5486 * C + 0.0043, (eq. 2) W = 1.3123 * C + -0.0016, and (eq. 3) E = -3.7319 * C + 0.1032. Roughly 15-20% of microorganisms reported to be anaerobes have amino acid compositions typical of aerobes.

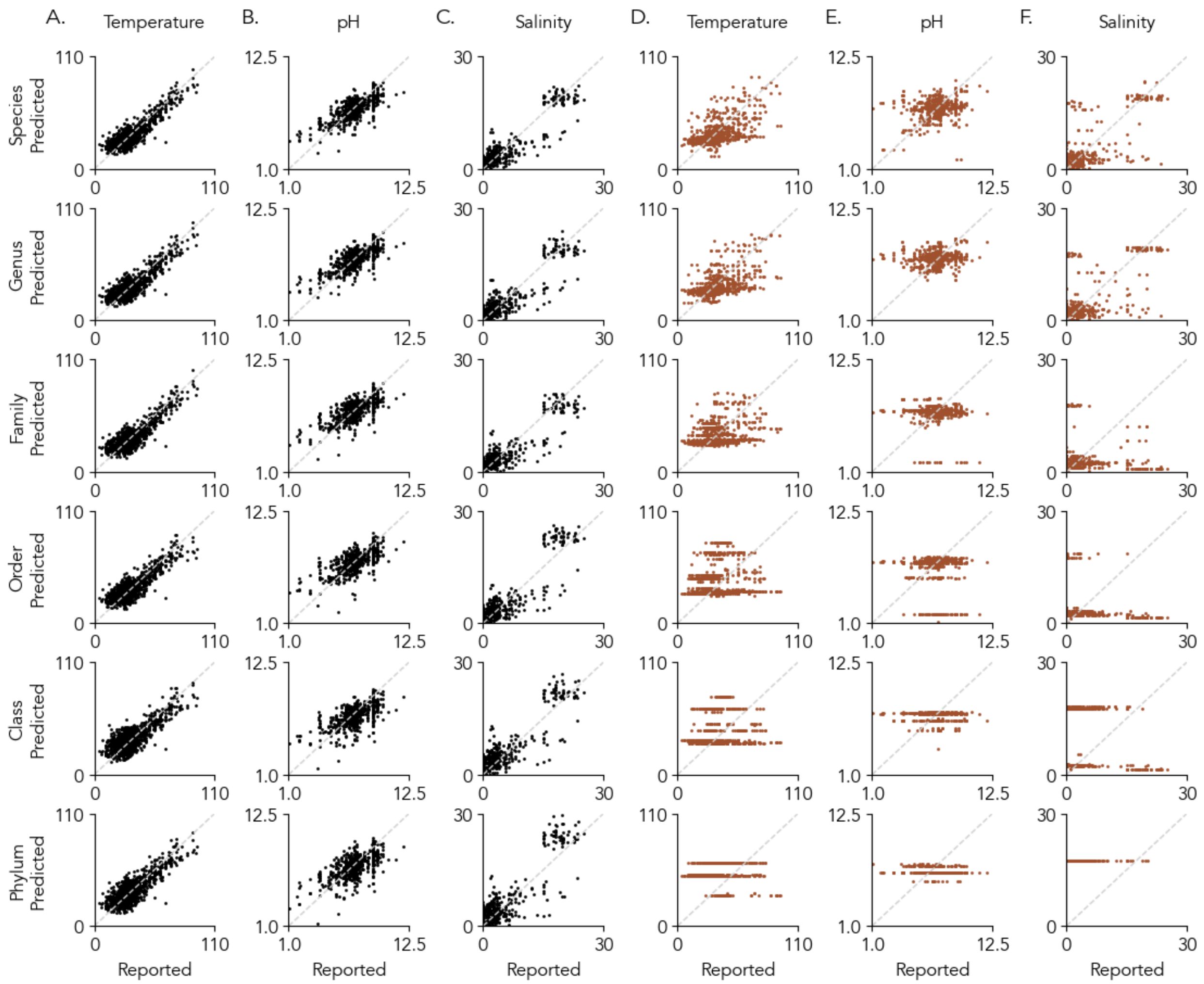


**Supplementary Figure 5. Holdout experiment scatter plots.**

Each plot compares reported (x-axis) and predicted (y-axis) values for the target condition of the column. Each row uses a different holdout level, ranging from species to phylum. The left set of plots (black) uses our models, and the right set of plots (red-brown) uses a simple phylogeny-based method where a "prediction" is the average value among the closest relatives of a species. As the holdout level increases, the difference between true and predicted values increases. Results are summarized in Figure 2C.

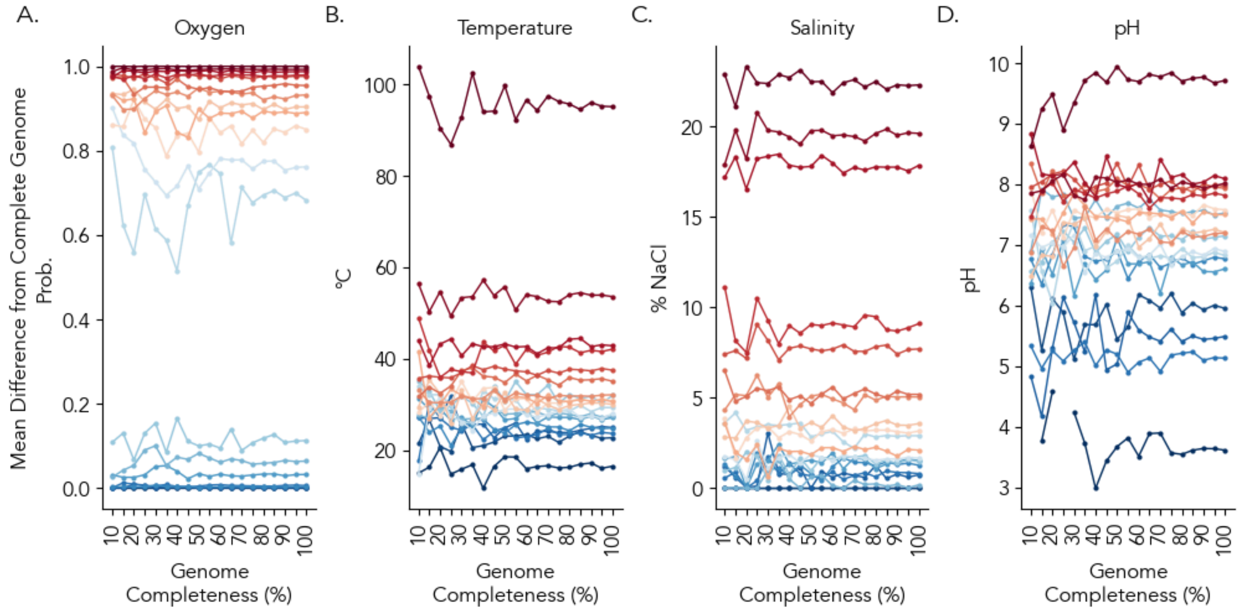


**Supplementary Figure 6. Accuracy vs. genome completeness for individual genomes.**

Twenty total genomes are shown, each genome is a different color. The y-axis shows the predicted value for a genome of a given percent completeness (x-axis). Genomes were selected to represent the typical distribution for each condition by selecting one genome at each of 20 percentiles. For example, few lines are above 40 C temperature because the vast majority of genomes in the dataset have moderate temperature preferences. Missing predictions (no line/points) are likely due to no random sampling of extracellular proteins, which are typically a small fraction of overall proteins yet required for several models.

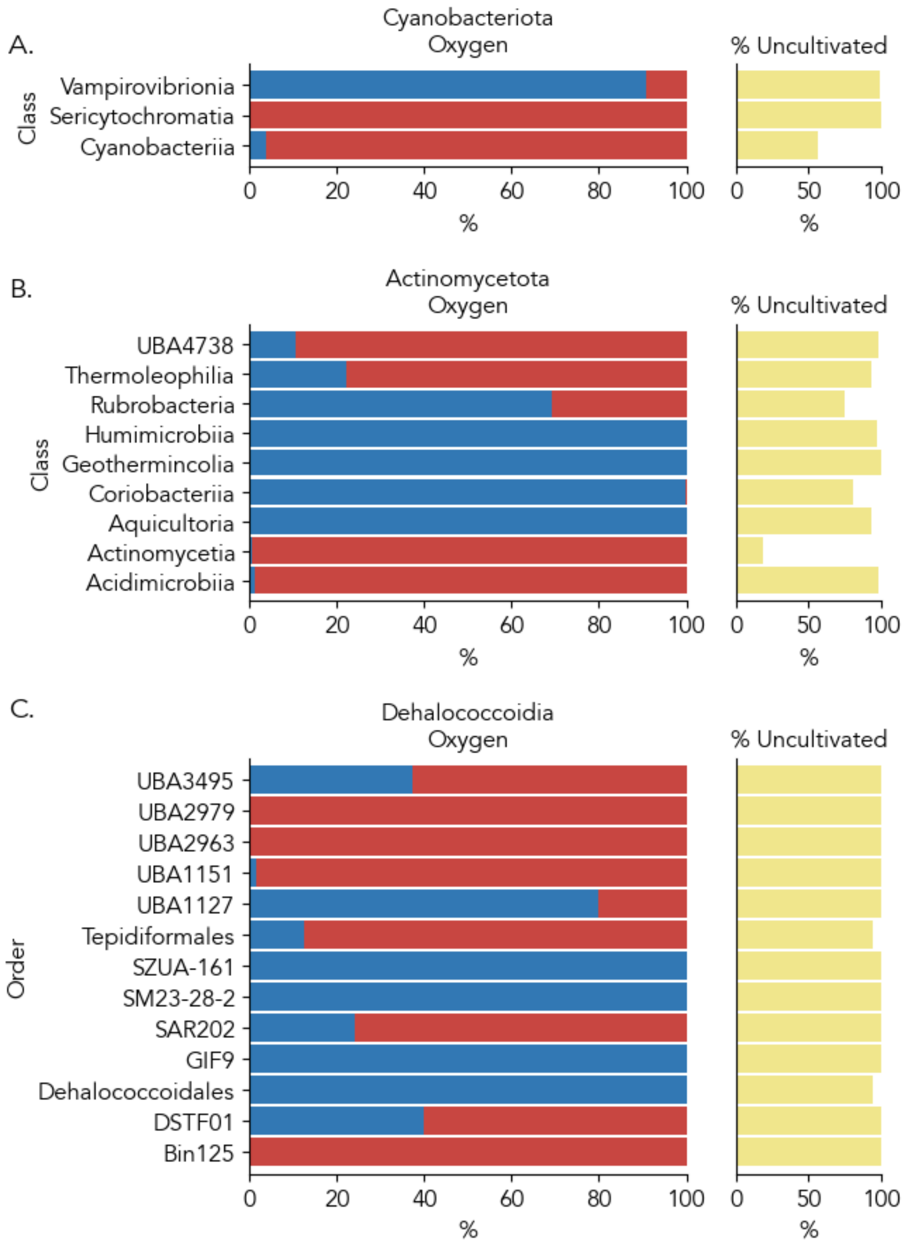


**Supplementary Figure 7. Examples of oxygen classifications for uncultivated lineages.**

Validation of the model by comparison to classes or orders of uncultivated microorganisms. Taxa with previously characterized oxygen tolerances are described in the main text. The percent of each taxon predicted to be oxygen intolerant (blue) or tolerant (red) are shown, and the % of species in the taxon that is uncultivated is indicated.

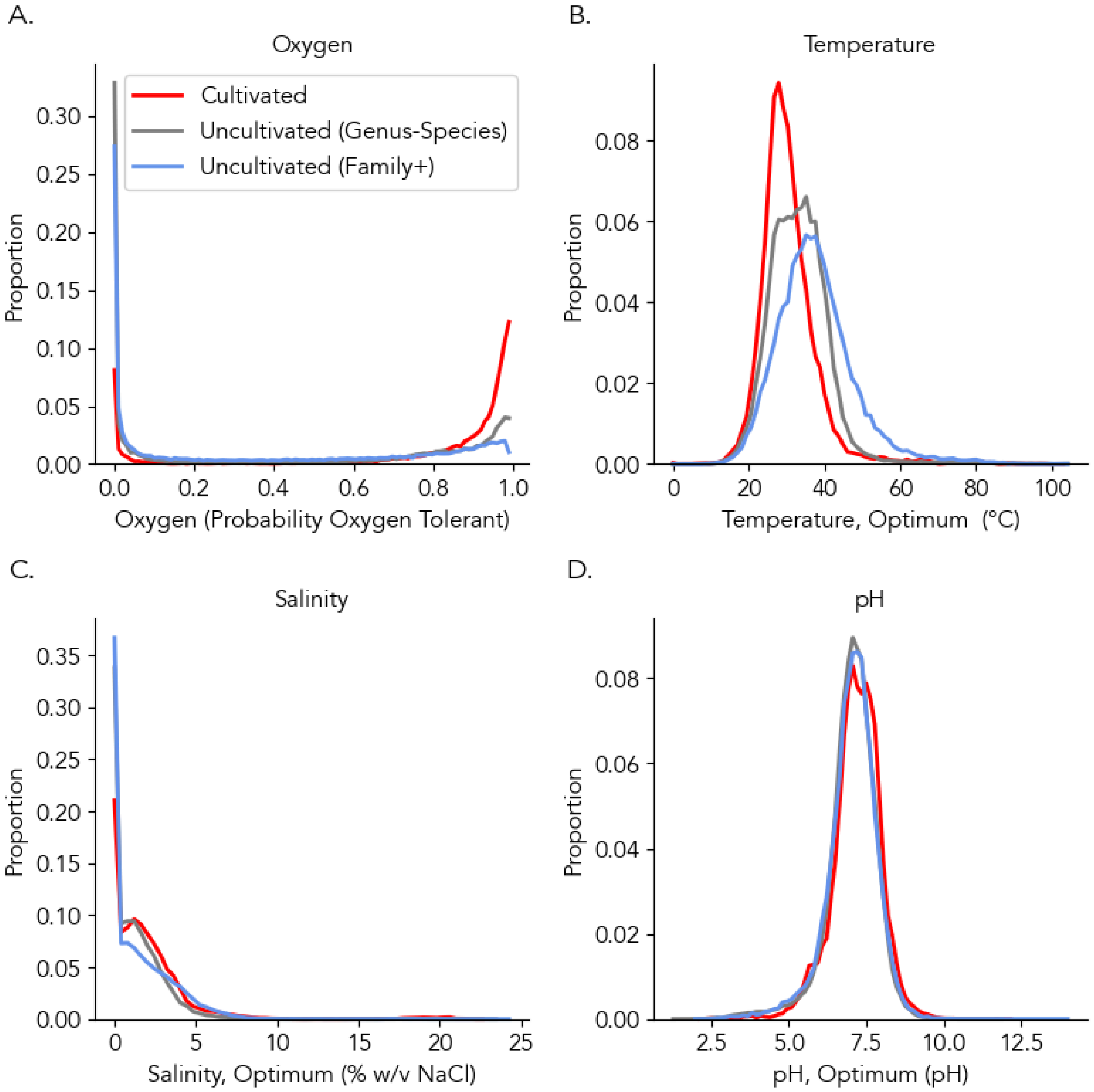


**Supplementary Figure 8. Trait distribution by cultivation status.**

Distribution of traits between cultivated species (red), uncultivated species within families with cultivated species (gray), and uncultivated species without cultivated members at the family level (blue).

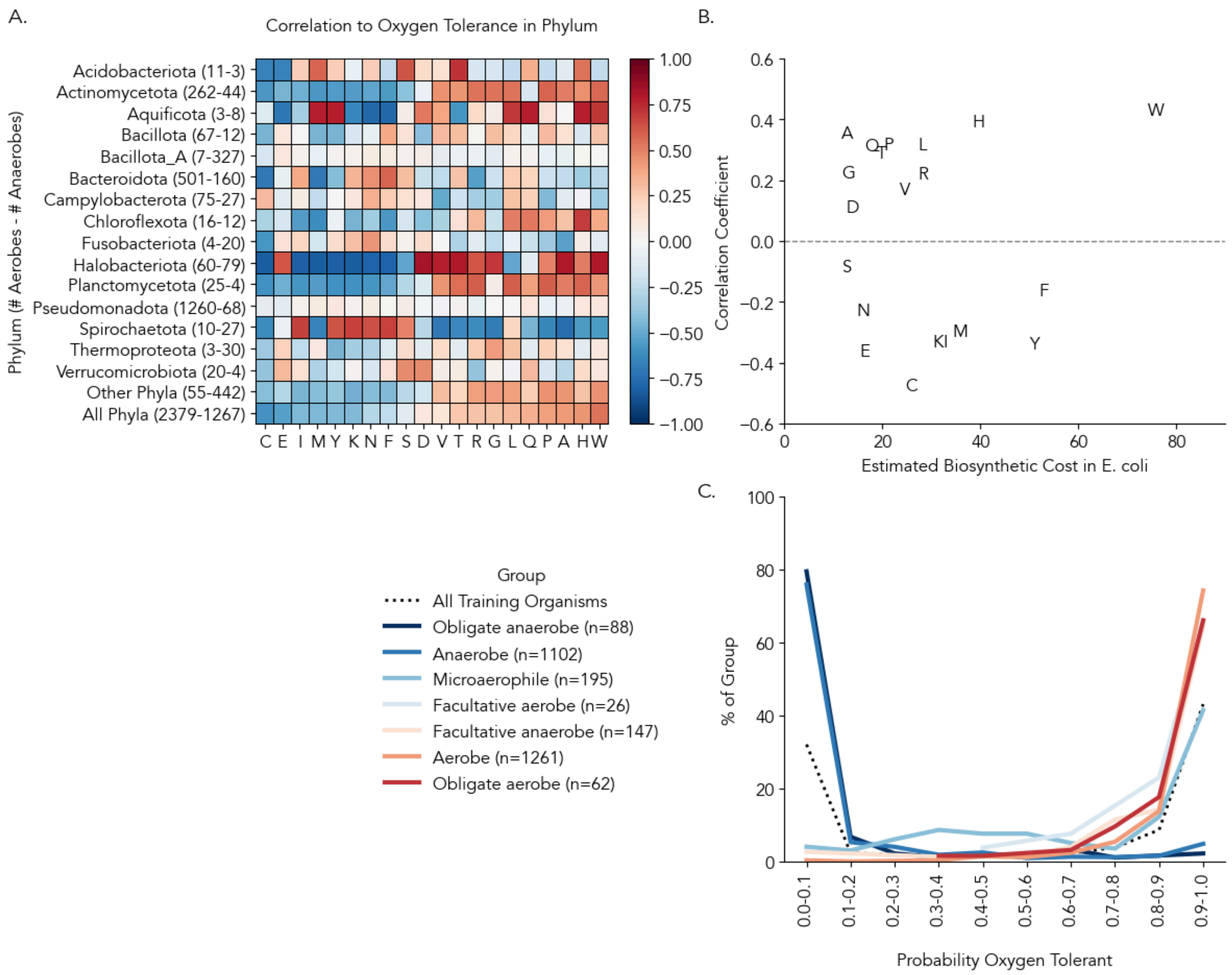


**Supplementary Figure 9. Understanding the oxygen model.**

(A) Heatmap showing the correlation of amino acid frequency to oxygen tolerance by phylum or groups of phyla. Values in parenthesis are the number of aerobes (left) and anaerobes (right) in a phylum, only showing phyla with 3+ of both. Color indicates the Spearman correlation coefficient. (B) Relationship between each amino acid's correlation across all phyla to oxygen tolerance (y-axis) and an estimate of the biosynthetic cost of amino acid synthesis in *E. coli* grown aerobically from Akashi et. al 2002 (x-axis). As described in the main text, amino acid biosynthesis costs will differ depending on the environment. (C) The distribution of predicted oxygen tolerance probability for different reported oxygen classifications. The x-axis is the predicted oxygen tolerance probability and the y-axis is the percent of the oxygen classification group found at that predicted probability. For example, 80% of organisms labeled "obligate anaerobes" are predicted to have a 0-0.1 chance of being oxygen tolerant. Note the relative rarity of predictions between 0.25-0.75 probability.

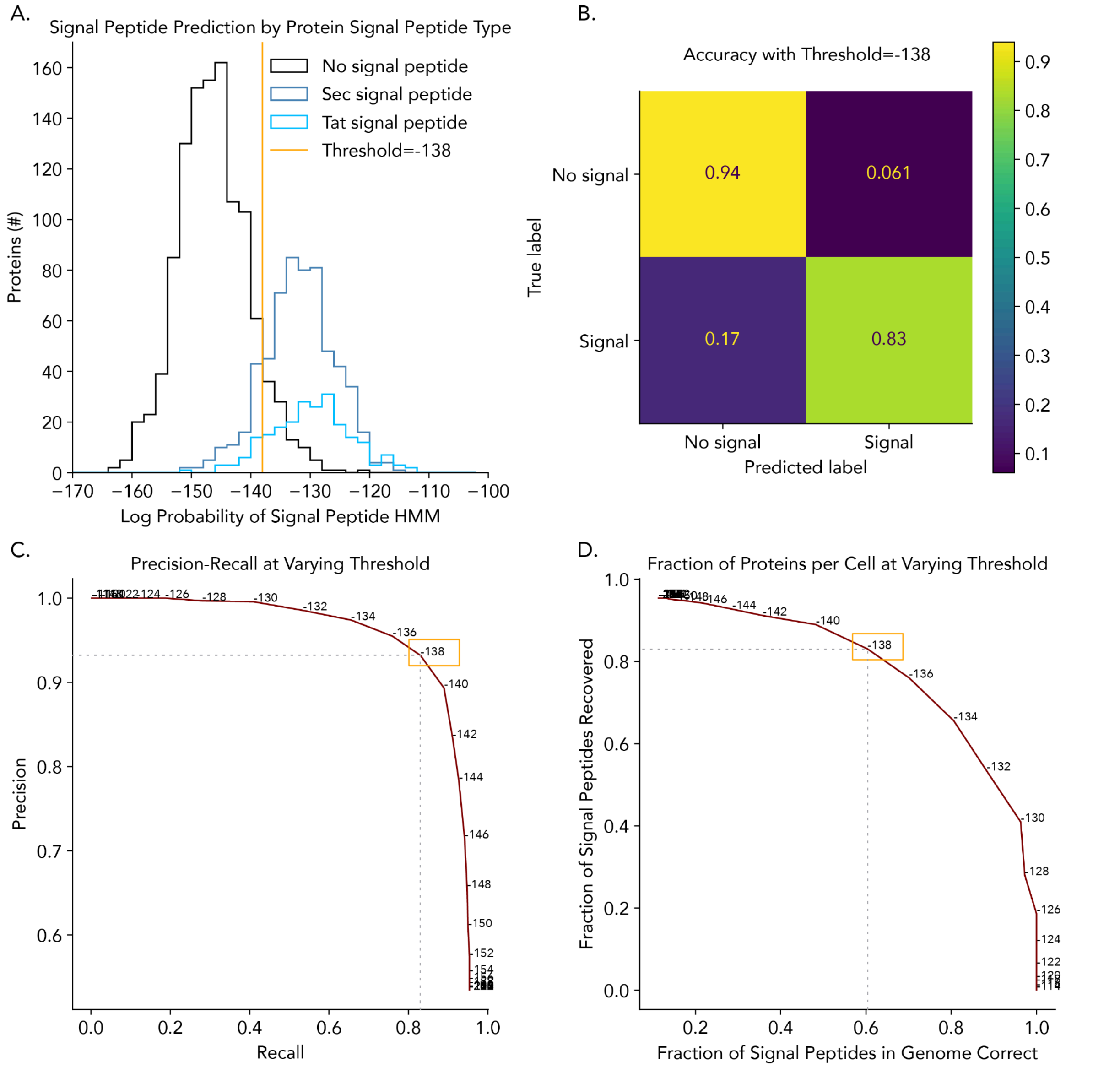


**Supplementary Figure 10. Construction of a hidden Markov model to predict signal peptides.**

The hidden Markov model assigns a relative probability of a Tat or Sec signal peptide to be present in the first 50 N-terminal amino acids. (A) The distribution of relative probability of being a signal peptide (x-axis) of proteins with or without different types of signal peptides (y-axis). The types of signal peptides are Tat (light blue), Sec (dark blue), and no signal peptide (black). The orange line is a threshold probability above which proteins are classified as having a signal peptide. (B) A confusion matrix indicating the accuracy of the above threshold to classify the training dataset: 94% of proteins without signal peptides and 83% of proteins with signal peptides were correctly classified. (C) A precision-recall curve showing how changing the threshold of classification affects the precision and recall. (D) A precision-recall curve as seen in (C) but in terms of the fraction of proteins in a genome, assuming only 10% of proteins in a genome are exported. Because 90% of proteins are not exported, false positive classifications can decrease the purity of proteins classified as exported. The chosen threshold value results in 20% of proteins to be false positives and captures 60% of exported proteins.

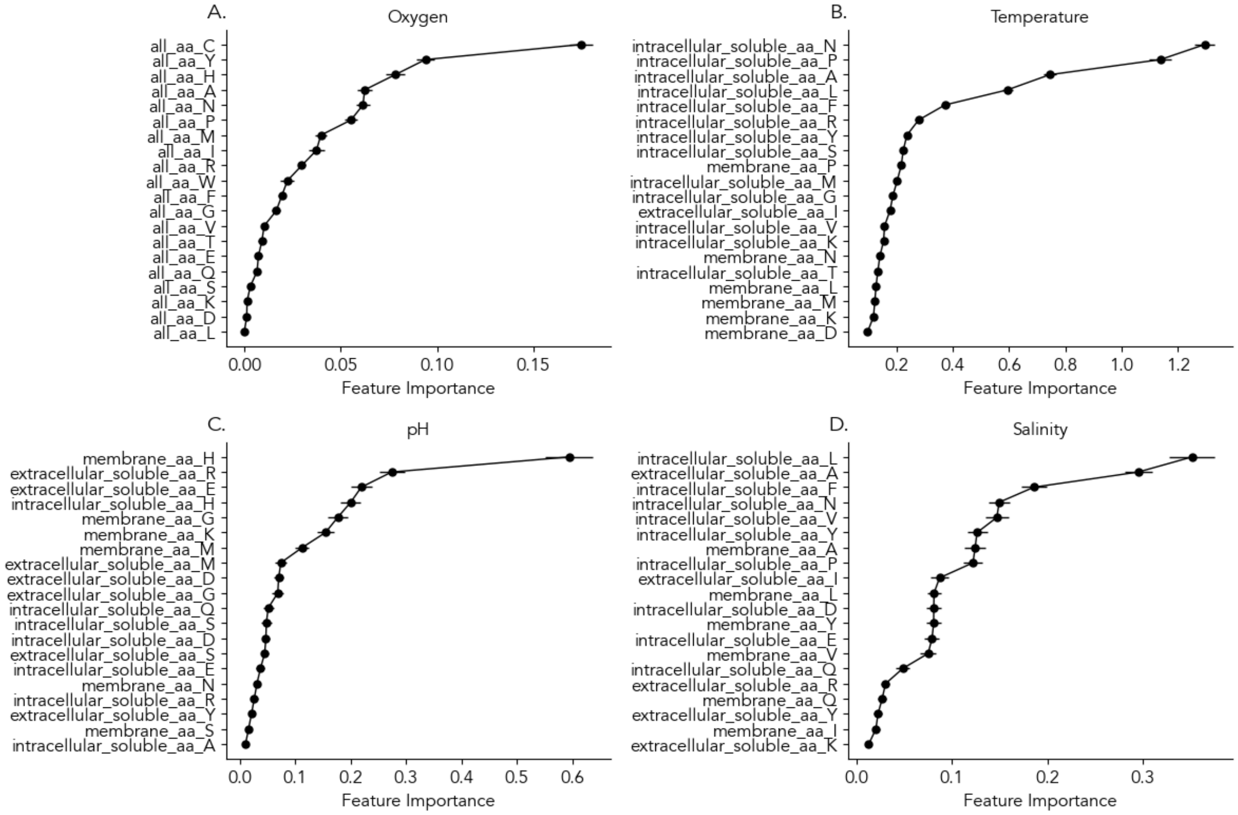


**Supplementary Figure 11. Feature importances in selected models for top 20 features.**

(A) Oxygen tolerance, (B) optimum temperature, (C) optimum salinity, and (D) optimum pH. Feature importances were calculated using permutation: measuring how much worse a prediction is when the values for a particular feature are shuffled across organisms. A higher feature importance score means that the feature is more important given a particular trained model. It does not necessarily mean that if the feature was not included, the model would not be as accurate.
